# Lysosomal dysfunction in Down Syndrome and Alzheimer mouse models is caused by selective v-ATPase inhibition by Tyr^682^ phosphorylated APP βCTF

**DOI:** 10.1101/2022.06.02.494546

**Authors:** Eunju Im, Ying Jiang, Philip Stavrides, Sandipkumar Darji, Hediye Erdjument-Bromage, Thomas A. Neubert, Jun Yong Choi, Jerzy Wegiel, Ju-Hyun Lee, Ralph A. Nixon

## Abstract

Lysosome dysfunction arises early and propels Alzheimer’s Disease (AD). Herein, we show that amyloid precursor protein (APP), linked to early-onset AD in Down Syndrome (DS), acts directly via its β-C-terminal fragment (βCTF) to disrupt lysosomal v-ATPase and acidification. In human DS fibroblasts, the phosphorylated ^682^YENPTY internalization motif of APP-βCTF binds selectively within a pocket of the v-ATPase V0a1 subunit cytoplasmic domain and competitively inhibits association of the V1 subcomplex of v-ATPase, thereby reducing its activity. Lowering APP-βCTF Tyr^682^ phosphorylation restores v-ATPase and lysosome function in DS fibroblasts and *in vivo* in brains of DS model mice. Notably, lowering APP-βCTF Tyr^682^ phosphorylation below normal constitutive levels boosts v-ATPase assembly and activity, suggesting that v-ATPase may also be modulated tonically by phospho-APP-βCTF. Elevated APP-βCTF Tyr^682^ phosphorylation in two mouse AD models similarly disrupts v-ATPase function. These findings offer new insight into the pathogenic mechanism underlying faulty lysosomes in all forms of AD.

## Introduction

The ultrastructural neuropathology of Alzheimer’s disease (AD) is dominated by robust and selective buildup of autophagic vacuoles within neurons, reflecting the defective transport and clearance of autophagic waste from neurons (*1, 2*). Abnormalities at all levels of the endosomal-lysosomal-autophagy network are an invariant early and progressive feature of AD neuropathology and pathophysiology (*1*). Accumulation of APP-βCTF in lysosomes is also believed to corrupt their functioning in mouse models of AD and Down Syndrome (DS, also referred to as Trisomy 21) and promote AD-related pathology (*3-5*).

The close association between lysosomal dysfunction and neurodegeneration is underscored by mutations of at least 30 genes operating within the endosomal-lysosomal system that cause inheritable neurodegenerative diseases across the aging spectrum (*2, 6*). In AD, disrupted lysosome function contributes to neuritic dystrophy, reduced clearance of β-amyloid and tau, synaptic plasticity deficits, and neurodegeneration (*7-10*). These deficits in mouse models are substantially remediated by restoring lysosomal functionality (*11-13*). Despite the growing evidence implicating lysosomes in pathogenesis in neurodegenerative diseases, the mechanisms underlying lysosomal system dysregulation remain unclarified.

Lumenal acidification of lysosomes to a pH of 4.5-5.0 is essential for their function (*14*), including optimal activation of varied hydrolytic enzymes and regulation of ion channels involved in lysosomal trafficking and cell signaling (*15, 16*). Lysosomal acidification is maintained primarily by a vacuolar (H^+^)-ATPase (v-ATPase), a multimeric enzyme complex that pumps protons from the cytosol into the lysosomal lumen (*14*). Fourteen protein subunits form the complete v-ATPase complex comprised of two sectors: an integral membrane-associated V0 sector and a cytosolic V1 sector capable of regulated dissociation from the V0 sector. The V1 sector, comprising subunits A-H, is responsible for ATP hydrolysis. The V0 sector, composed of subunits a, d, e, c, and c’’, forms the channel that conducts proton transport (*17, 18*).

v-ATPase activity is subject to various forms of regulation that modulate subunit expression, assembly, trafficking, and signaling (*18*). Among the most important is regulation of the V1 sector association with the membrane-bound V0 sector (*19, 20*). Cryogenic electron microscopy analysis showed that ATP hydrolysis in the soluble catalytic region (V1 sector) of v-ATPase is coupled to proton translocation through the membrane-bound region (V0 sector) by rotation of a central rotor subcomplex, with peripheral stalks preventing the entire membrane-bound region from turning with the rotor (*21*). Thus, to form a functional channel of v-ATPase, assembly of V0 sector and V1 sector is a necessary event. Physiological changes in v-ATPase activity modulate nutrient sensing that controls mTOR activity, autophagy induction, and lysosomal biogenesis (*15, 22, 23*). Mutations of v-ATPase subunits cause nearly a dozen familial degenerative diseases, most having a significant CNS phenotype (*24*). v-ATPase components are most highly expressed in neurons and activity is regulated by various signaling cascades that likely contribute to the way different risk factors compromise lysosomes in AD (*24*). Notably, we discovered that familial early-onset AD due to *PSEN1* mutation disrupts maturation and lysosomal delivery of v-ATPase V0a1 subunit, thus impairing v-ATPase assembly and proton pumping and leading to diverse features of AD pathophysiology (*25*).

In this study, we show that *APP*, like *PSEN1* in AD, disrupts v-ATPase complex assembly and lysosomal acidification by a distinctive mechanism that also directly involves the V0a1 subunit. We demonstrate that phospho-Tyr^682^-APP-βCTF (pY^682^APP-βCTF) generated by β-secretase (BACE-1) cleavage from constitutively phosphorylated APP, selectively and directly interacts with the cytoplasmic domain of the V0a1 subunit. This interaction impedes association of V1 subunits thereby inhibiting v-ATPase activity and acidification of lysosomes. Levels of pY^682^APP-βCTF can rise by multiple mechanisms, including elevated APP expression, increased BACE1 activity, under-active protein phosphatases, or an over-active Tyr kinase. Regardless of which combination of the foregoing mechanisms operates in DS or AD, we show that lowering APP Tyr^682^ phosphorylation pharmacologically can prevent the disruptive effects of APP-βCTF on lysosome function *in vivo*. Our results and growing knowledge of AD pathobiology support our view that v-ATPase disruption is a primary triggering mechanism in AD and that v-ATPase dysregulation is a new therapeutic target. Finally, our initial evidence that lowering APP levels or APP-βCTF itself below baseline endogenous levels accentuates v-ATPase assembly and activity, raises the possibility of tonic modulation of lysosomal function by APP in physiological states as well as in disease. Our findings reported here and other emerging data (*8, 26*) point toward a unifying pathogenic mechanism underlying primary lysosomal dysfunction in AD.

## Results

### Lysosomal v-ATPase dysfunction in DS fibroblasts is caused by impaired complex assembly

We confirmed that lysosomal acidification is impaired in Down Syndrome (DS) fibroblasts (*5*) by measuring lysosomal pH using LysoSensor™ Yellow/Blue dextran, a ratiometric dye targeted to lysosomes (*27*). Lysosomal pH was significantly increased in DS fibroblasts compared to control (2N) (Fig. 1A). Since lysosome acidity is generated mainly by v-ATPase (*14*), we measured lysosomal v-ATPase activity by using a superparamagnetic chromatography isolation procedure (*28*) to isolate highly enriched lysosomal preparations from DS and 2N cells, the purity of which is reflected by the high level of LAMP1 enrichment (fig. S1). We then measured the rate of ATP hydrolysis in purified lysosomes monitored by release of inorganic phosphate, which revealed ∼40% lower rates in lysosomes isolated from DS fibroblasts relative to 2N (Fig. 1B). To confirm the predicted impact of this lowered ATP hydrolysis, we measured the rates of proton translocation into the lysosomal lumen using the quenching of 9-amino-6-chloro-2-methoxyacridine (ACMA), which uncovered a ∼80% lower rate in purified lysosomes from DS fibroblasts than from 2N lysosomes (Fig. 1, C to D).

**Fig. 1.**
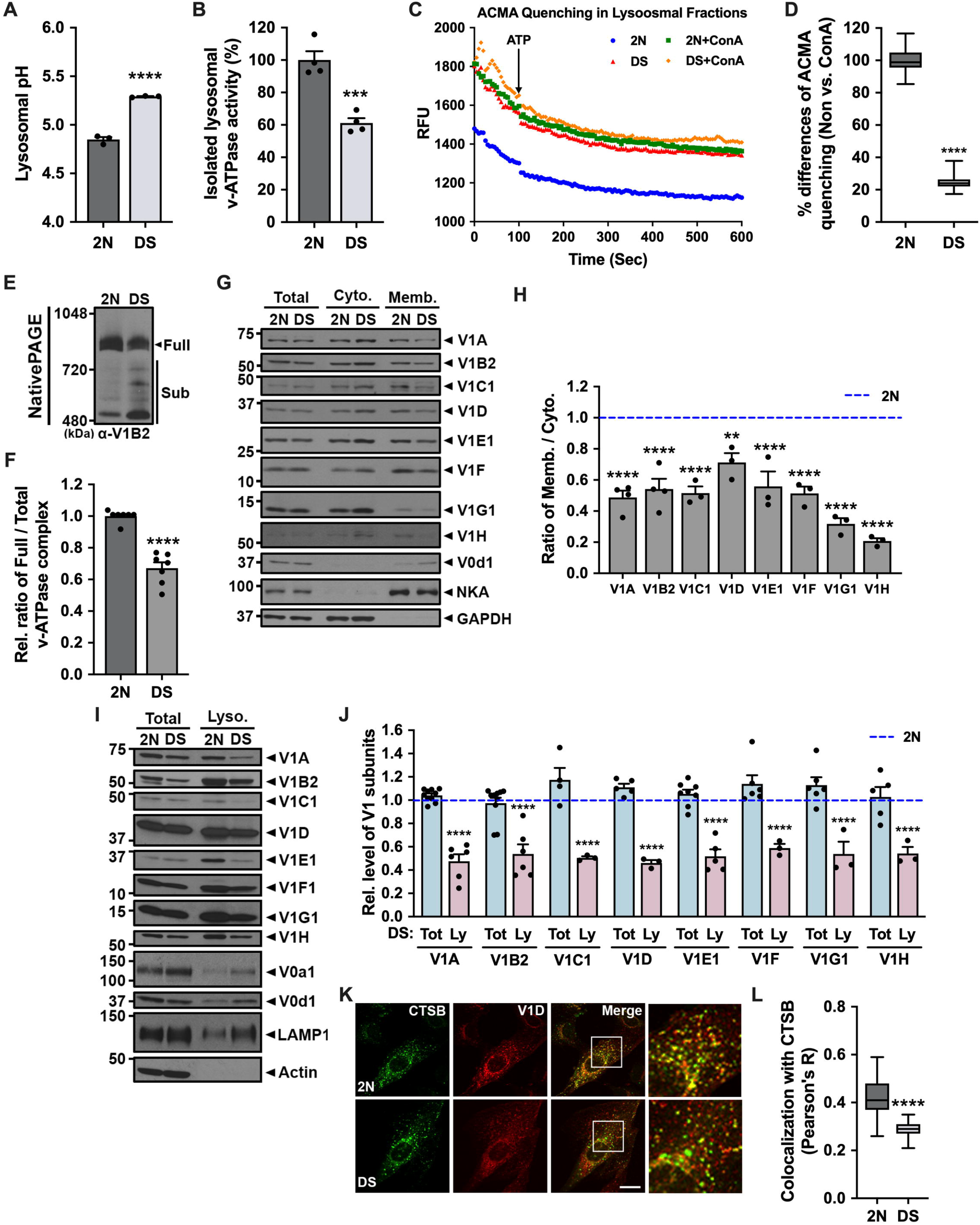
Assembly of the lysosomal v-ATPase complex is impaired in DS fibroblasts. **(A)** Lysosomal pH of 2yr 2N and DS fibroblasts measured ratiometrically using LysoSensor Yellow/Blue (Y/B) dextran (n=3, three independent, triplicate). **(B-D)** Lysosomal v-ATPase activity of 2yr 2N and DS fibroblasts measured colorimetrically as ATP hydrolysis (*B*) (n=4, four independent, duplicate) and fluorometrically (ACMA method) as H^+^ transport (*C, D*) (n=3, three independent) using Lyso fractions. **(E)** Membrane fractions of 2yr 2N and DS fibroblasts were resolved using native PAGE and immunoblotted with anti-V1B2 antibody. **(F)** The graph represents relative ratio of full complex divided by total (full plus sub complexes) complexes of v-ATPase (n=7, seven independent). **(G)** Immunoblots of v-ATPase subunits distribution in cytosol (Cyto.) and membrane (Memb.) fractions of 2yr 2N and DS fibroblasts. Na, K-ATPase 1 (NKA) served as a Memb. maker, and GAPDH served as a Cyto. marker. **(H)** The graphs represent ratio of Memb. vs Cyto. band of each v-ATPase subunits (n≥3, at least three independent). **(I)** Immunoblot of v-ATPase subunit distributions in total lysates (Total) and lysosome enriched (Lyso.) fractions of 2yr 2N and DS fibroblasts. LAMP1 served as a marker for lysosomes and Actin served as a loading control for total lysates. **(J)** The graphs show relative band intensity for each v-ATPase subunit (n≥3, at least three independent). **(K, L)** Double-immunofluorescence labeling in 2yr 2N and DS fibroblasts shows colocalization of v-ATPase (V1D) and cathepsin B (CTSB) (*K*). Scale bar, 10 µm. Quantification analysis of v-ATPase V1D and lysosomal marker, CTSB, shows colocalization in 2N and DS fibroblasts as calculated by Pearson’s correlation coefficient (n ≥ 131 cells, three independent) (*L*). Quantitative data (A, B, D, F, H, J, and L) are presented as mean values with ±SEM, two-tailed unpaired t test (A, B, D, F, and L), ordinary one-way ANOVA with Šidák’s multiple comparisons test (H), ordinary two-way ANOVA with Šidák’s multiple comparisons test (J); **, P < 0.005; ***, P < 0.0005; ****, P < 0.0001. Each dot represents average value of technical replicates from each independent experiment.

The v-ATPase complex is only active when the membrane-bound V0 sector and the cytosolic V1 sector are assembled on the lysosomal membrane (*20*). We examined the assembly of the v-ATPase complex using native polyacrylamide gel electrophoresis (PAGE) followed by mass spectrometric (MS) analysis. On native PAGE gels, the fully assembled v-ATPase (about 880 kDa) was present at significantly reduced in DS fibroblasts as compared to 2N cells (Fig. 1E; “Full”). Furthermore, DS cells contained elevated levels of incompletely assembled subcomplexes exhibiting increasing mobility on the native gel, which correlated with detection of fewer V1 subunits associated with V0 in the membrane fraction (Fig. 1E; “Sub”). The relative ratio of fully assembled v-ATPase divided by total v-ATPase (fully assembled v-ATPase plus subcomplexes of v-ATPase bands) revealed ∼35% lower ratios in DS fibroblasts as compared to 2N (Fig. 1F). This indicates that decreased lysosomal v-ATPase activity was due to reduced levels of the fully assembled functional v-ATPase complex formation in DS fibroblasts. The bands of v-ATPase were detected with a V1B2 subunit antibody and shown by MS analysis to contain all V1 subunits (fig. S2, A to B). To further investigate the possible disruption of v-ATPase assembly in DS cells, we examined v-ATPase complex assembly on cell membranes separated from the cytosol, each subjected to immunoblot analysis. Because affinities of the specific antibodies differ for each subunit, the relative abundance of subunits cannot be compared by band intensity; however, the membrane (Memb):cytosol (Cyto) ratio for each subunit is a reliable index of the changes in degree of association of a given V1 subunit with the V0 sector. Importantly, the Memb:Cyto ratios for all of the V1 subunits were significantly lowered in DS fibroblasts (Fig. 1, G to H).

Because APP-βCTF are implicated in lysosomal dysfunction (*3, 5*) and accumulate in lysosome-related compartments in AD and DS models, we next investigated v-ATPase assembly and function using lysosome enriched (Lyso) fractions, which indeed demonstrated a disproportionate v-ATPase deficit relative to the total cellular v-ATPase pool. All V1 subunits in purified lysosomes from DS fibroblasts were present at markedly decreased levels compared to those in the Lyso fraction from 2N cells (Fig. 1, I to J). Importantly, these findings on isolated lysosomes were confirmed by further results from double-immunofluorescence labeling analyses that revealed strong colocalization of V1D subunit with cathepsin B (CTSB)-positive compartments in 2N cells (Fig. 1K). Quantitative analyses of colocalization with lysosomes showed a significantly decreased V1D level in DS fibroblasts (Fig. 1L). We confirmed lysosomal pH, v-ATPase activity and assembly, and localization of V1 subunits in another line of DS fibroblasts (5 month age) compared to age matched control (2N) fibroblasts (fig. S3).

Taken together, the foregoing data indicated that dysfunction of v-ATPase activity is related to a markedly lowered association of the V1 sector with the V0 sector on lysosomal membranes in DS fibroblasts.

### APP-βCTF modulates v-ATPase activity and is inhibitory at elevated levels in DS

In light of previous studies linking βCTF to lysosomal dysfunction in AD, DS, and AD models (*3, 5, 8*), we investigated whether the mechanism of the underlying lysosomal v-ATPase deficits in DS fibroblasts was due to elevated generation of APP-βCTF from the extra APP gene copy (3N). We knocked down APP expression to levels below those in 2N control cells using small interference RNA constructs specifically against human *APP* (siAPP) and compared these cells to those exposed to scrambled RNA sequences (siNC). Knock-down (KD) of APP, confirmed by western blot (WB) 72 hr after siRNA transfection, significantly rescued lysosomal pH (Fig. 2A) and v-ATPase activity (Fig. 2B) in DS fibroblasts. Moreover, the same treatment restored assembly of the v-ATPase complex (Fig. 2, C to D) with a full complement of V1 subunits (Fig. 2, E to F) in DS fibroblasts. These data therefore indicate that, even in the presence of the entire extra copy of Chr.21 in DS cells, v-ATPase dysfunction is dependent mainly on the extra copy of APP. Unexpectedly, knockdown of APP in 2N fibroblasts further lowered lysosomal pH by 0.11 pH units which was associated with significantly higher v-ATPase activity (16%). Moreover, levels of V1 subunits associated with lysosomes were significantly higher while corresponding levels in the cytosol were lowered, consistent with an increased assembly of v-ATPase complex on lysosomes in 2N control fibroblasts with APP levels below normal baseline. These findings in 2N fibroblasts and additional data discussed later suggest that v-ATPase may be tonically modulated by APP.

**Fig. 2.**
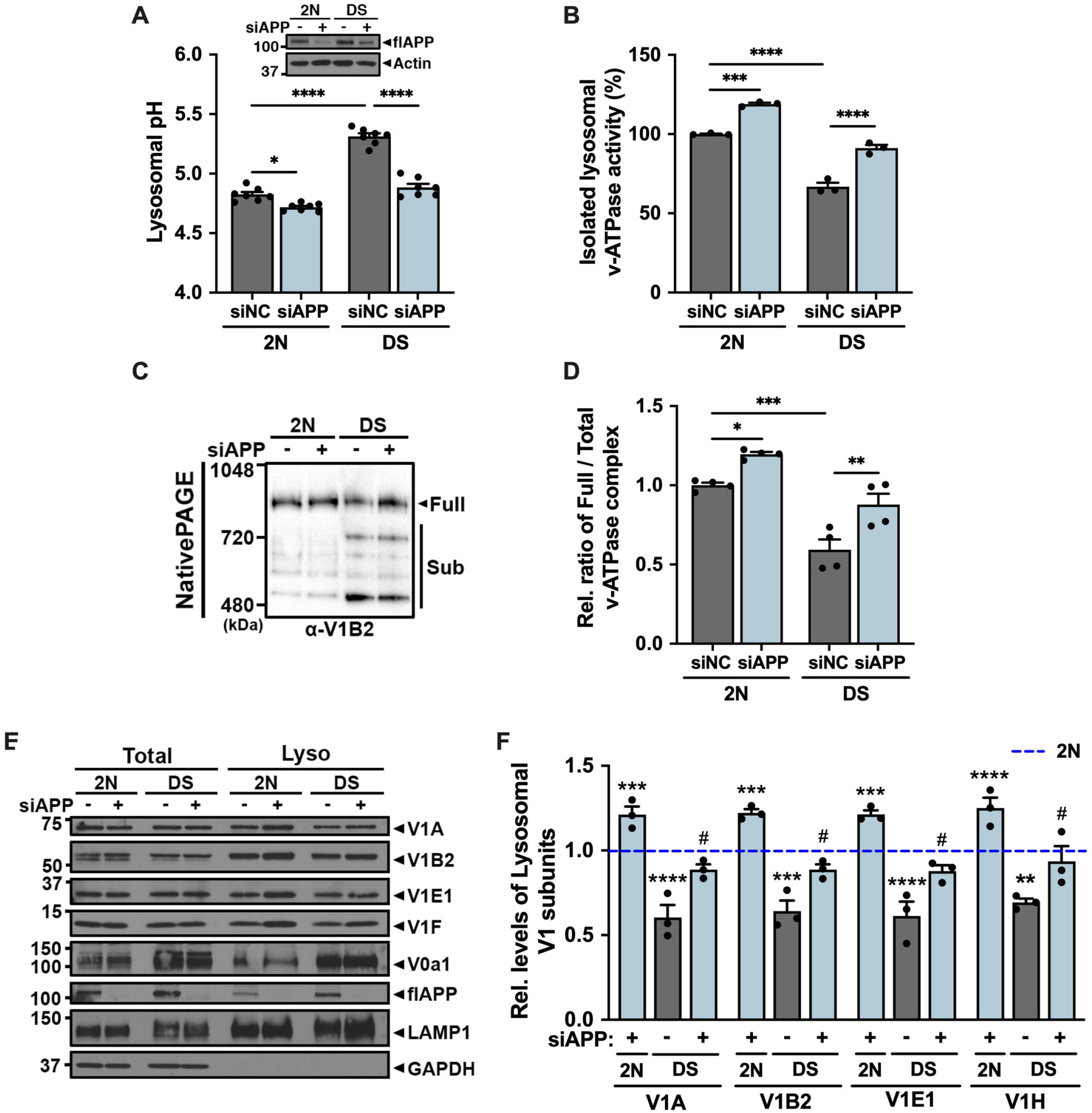
Impairment of lysosomal v-ATPase in DS fibroblasts is dependent on APP expression. **(A)** Lysosomal pH of 2yr 2N and DS fibroblasts transfected with siRNA for either negative control (siNC) or APP (siAPP) for 72 hr determined by LysoSensor Y/B dextran (n=7, seven independent, quadruplicate). The immunoblot represents APP levels in 2N and DS fibroblasts after transfection with 40 nM siNC or siAPP. **(B)** ATP hydrolysis activity of v-ATPase measured using lysosomal fractions from siRNA transfected 2yr 2N and DS fibroblasts (n=3, three independent, duplicate). **(C)** Membrane fractions of siRNA transfected 2yr 2N and DS fibroblasts were resolved using native PAGE and immunoblotted with anti-V1B2 antibody. **(D)** The graph represents relative ratio of full complex divided by total (full plus sub complexes) complexes of v-ATPase (n=4, four independent). **(E)** Immunoblots of v-ATPase subunit distributions in total and Lyso. fractions of siRNA transfected 2yr 2N and DS fibroblasts. LAMP1 served as a Lyso marker and GAPDH served as a loading control for total lysates. **(F)** The graphs show band intensity of each v-ATPase subunit from Lyso fraction (n=3, three independent). Quantitative data (A, B, D, and F) are presented as mean values with ±SEM, ordinary one-way ANOVA with Šidák’s multiple comparisons test (A, B, and D), ordinary two-way ANOVA with Šidák’s multiple comparisons test (F); *, P < 0.05; **, P < 0.005; ***, P < 0.0005; ****, P < 0.0001. Statistical significance between groups is shown by symbols: *2N+siNC vs others, #DS+siNC vs DS+siAPP. Each dot represents average value of technical replicates from each independent experiment.

APP-βCTF levels are elevated in AD and DS brain (*29, 30*) and rise substantially in lysosomes (*3, 5*). We observed that such increases in DS fibroblasts (*5*) are accompanied by lysosomal pH elevation (Fig. 3A). We addressed the specificity of APP-βCTF as a mediator of this effect by first measuring v-ATPase activity in lysosomes from cells treated with vehicle or γ-secretase inhibitor L685,458 (γ-Sec INH), which elevates APP-βCTF levels (fig. S4A) by inhibiting its γ-cleavage to form Aβ peptide (*31*). In 2N cells, γ-Sec INH elevated APP-βCTF levels and strongly inhibited v-ATPase activity (Fig. 3B). In DS fibroblasts where APP-βCTF is already elevated and v-ATPase activity is disrupted, an additional impact of γ-Sec INH was not detected (Fig. 3B). We observed this same pattern of DS phenotype induction in 2N cells when assembly of lysosomal v-ATPase was assessed by native gel analysis (Fig. 3, C to D) or CTSB and V1D colocalization was assayed by double-label IFC (Fig.3, E to F).

**Fig. 3.**
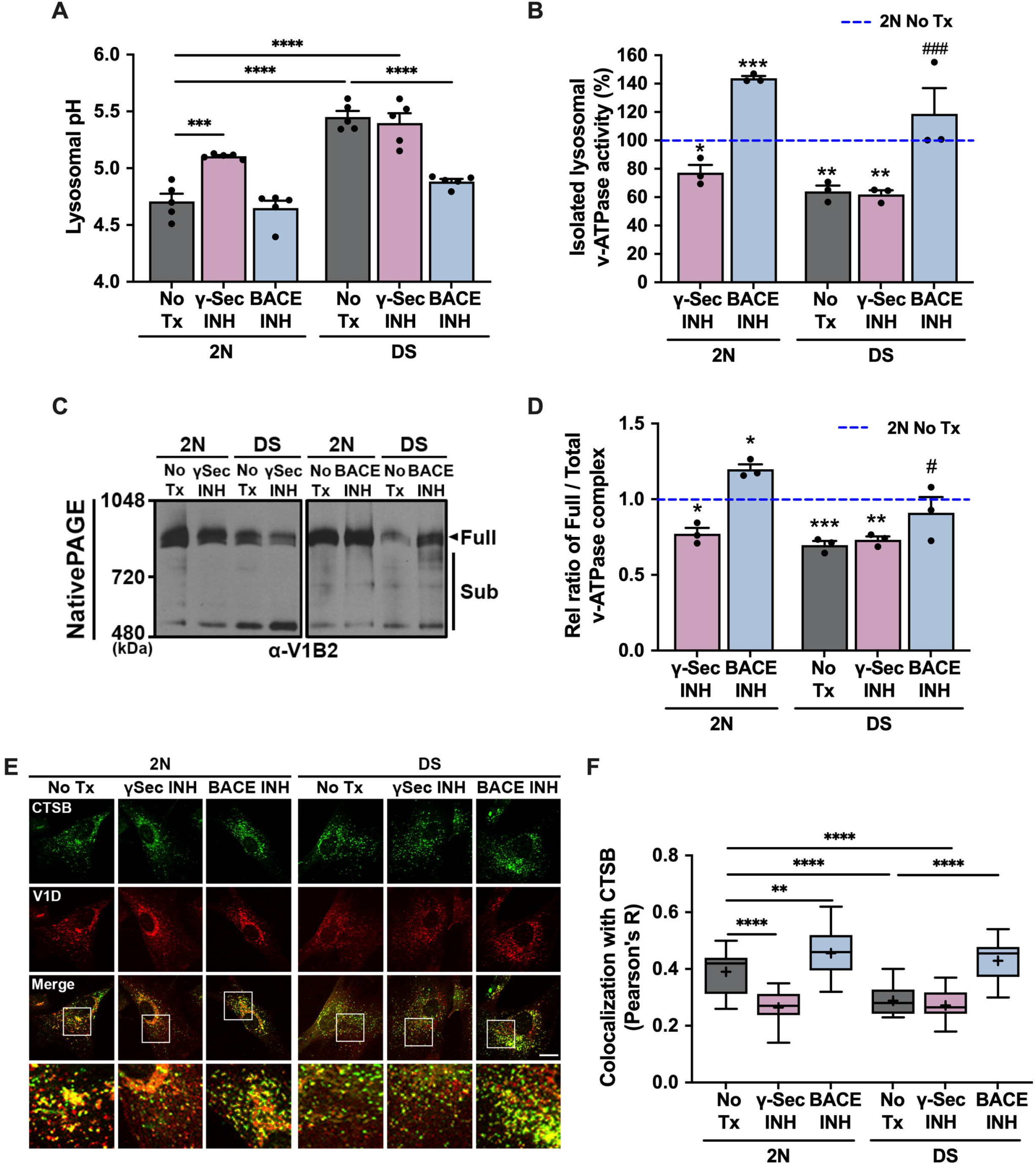
Lysosomal v-ATPase activity is specifically affected by APP-βCTF among metabolites of APP. **(A)** Lysosomal pH of 2yr 2N and DS fibroblasts treated with either DMSO (No Tx), γ-secretase inhibitor, L685,458 (γ-Sec INH; 10 µM), or BACE1 inhibitor (BACE INH; 10 µM) for 24 hr determined by LysoSensor Y/B dextran (n=5, five independent, triplicate). **(B)** ATP hydrolysis activity of v-ATPase measured using lysosome fractions from 2yr 2N and DS fibroblasts after treatment indicated inhibitors (n=3, three independent, duplicate). **(C)** Membrane fractions from 2yr 2N and DS fibroblasts treated with indicated inhibitors were resolved using the native PAGE and immunoblotted with anti-V1B2 antibody. **(D)** The graph represents relative ratio of full complex divided by total (full plus sub complexes) complexes of v-ATPase (n=3, three independent). **(E, F)** Double-immunofluorescence labeling shows colocalization of V1D and CTSB in 2N and DS fibroblasts treated with indicated inhibitors (*E*). Scale bar, 10 µm. Quantification analysis of v-ATPase V1D and lysosomal marker, CTSB, shows colocalization in 2N and DS fibroblasts treated with indicated inhibitors as calculated by Pearson’s correlation coefficient (n ≥ 123 cells, three independent) (*F*). Quantitative data (A, B, D, and F) are presented as mean values with ±SEM, ordinary one-way ANOVA with Šidák’s multiple comparisons test; *, P < 0.05; **, P < 0.005; ***, P < 0.0005; ****, P < 0.0001. Statistically significance between groups is shown by symbols: *2N No Tx vs others, #DS No Tx vs DS+inhibitor. Each dot represents average value of technical replicates from each independent experiment.

In contrast to effects of γ-Sec INH, lowering APP-βCTF generation from APP by using BACE1 inhibitor IV (BACE INH) (fig. S4B) almost fully recovered lysosomal pH (Fig. 3A) and function of v-ATPase in DS fibroblasts (Fig. 3, B to F). Further supporting the concept that APP may tonically modulate v-ATPase activity, v-ATPase function in 2N fibroblasts was enhanced by BACE INH, similar to observations in the siAPP studies (Fig. 2B), ATPase activity rose significantly (50%) above baseline activity in BACE INH-treated 2N cells (Fig. 3B) associated with significantly enhanced v-ATPase complex assembly (Fig. 3, C to D) and V1D colocalization with CTSB-positive lysosomes (Fig. 3, E to F). Furthermore, lysosomal pH trended non-significantly toward a lower than baseline level (Fig. 3A). Finally, to rule out an effect of APP-⍰CTF on v-ATPase function, we blocked its generation in cells using the selective ⍰-secretase inhibitor TAPI-1 (*32*), which reduced ⍰CTF levels by 30-50% (fig. S5A). No discernible changes of lysosomal pH (fig. S5B), ATPase activity (fig. S5C) or v-ATPase assembly (fig. S5, D to E) were seen in 2N and DS fibroblasts after exposure to TAPI-1.

Taken together, the foregoing data indicated that v-ATPase is specifically regulated by APP-βCTF and suggested that the baseline level of v-ATPase function in 2N fibroblasts reflects a tonic balance that can be potentially modulated by APP-βCTF levels.

### APP-βCTF binds selectively to v-ATPase via its YENPTY domain and depends on Tyr^682^ APP phosphorylation

A previous synaptic interactome analysis of the APP intracellular domain (AICD) revealed APP-interacting proteins associated with pre-synaptic vesicle synaptic termini including several v-ATPase subunits, such as V0a1, V0d1, V1A, V1B2, V1C, V1D, and V1E (*33*). Based on this clue, we investigated whether APP interacts with v-ATPase subunits by performing co-immunoprecipitation (IP) analyses in 2N and DS fibroblasts. In control 2N cell lysates, V0a1, V0d1, V1A, V1B2, V1D, and V1E1 were pulled down in IP using C1/6.1 antibody against the extreme C-terminal sequence of APP (amino-acid residues 676-695 of human APP 695 isoform) but not in immune-precipitates using a non-specific control antibody (IgG) (Fig. 4, A to B, left panel). The binding of APP with V1 subunits (V1A, V1B2, V1D, and V1E1) was decreased in DS cell lysates (Fig. 4B, left panel), while APP strongly interacted with V0 subunits (V0a1 and V0d1) (Fig. 4A, left panel). To establish a lysosomal localization of the interaction between APP and v-ATPase subunits, we performed co-IP assays using a Lyso fraction. APP showed elevated interaction with V0a1/V0d1 in the Lyso fraction of DS fibroblasts (Fig. 4C). By contrast, APP did not interact with V1 subunits (V1A, V1B2, V1D, and V1E1) in the Lyso fraction (Fig. 4C).

**Fig. 4.**
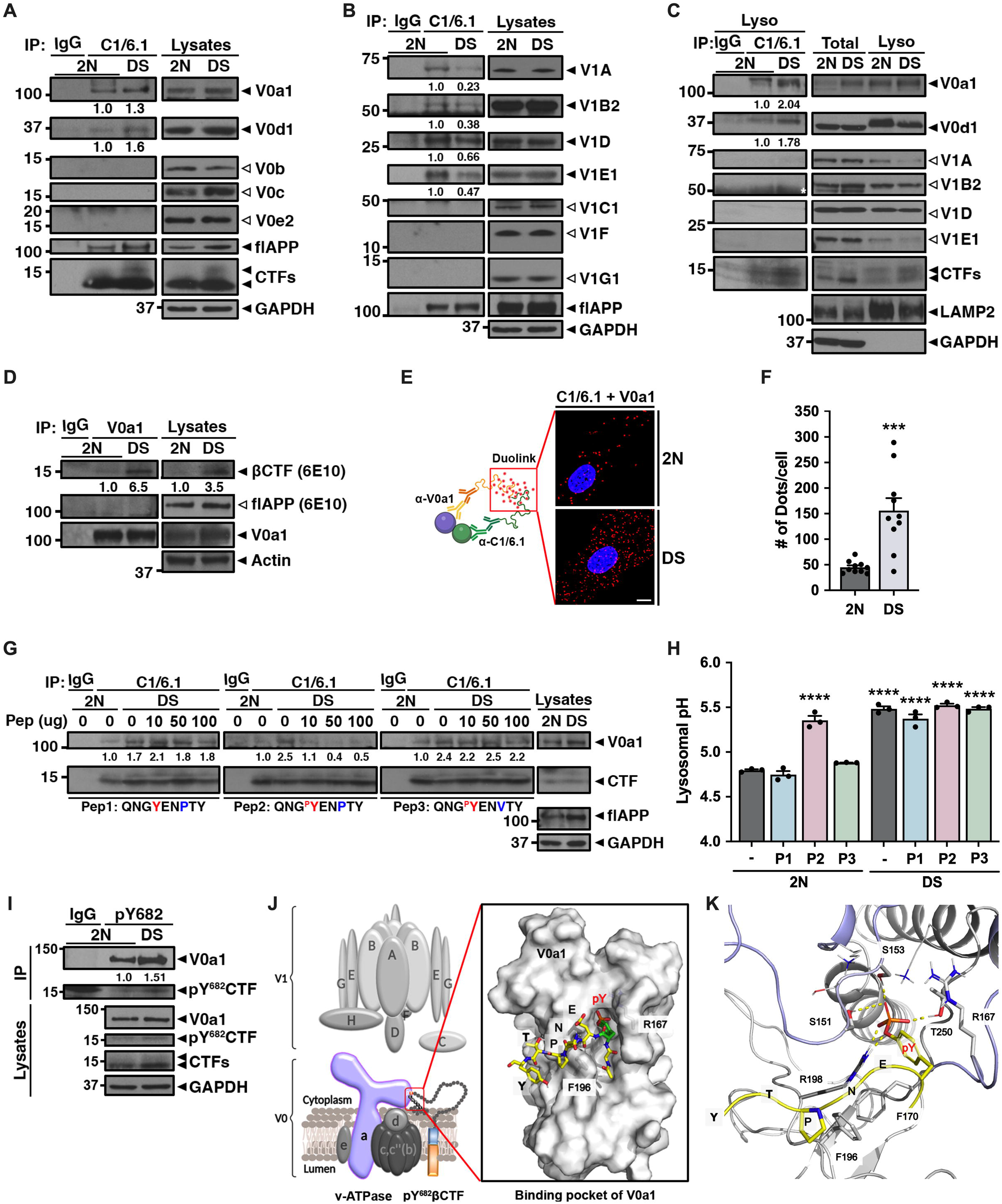
pYENPTY motif of APP-βCTF interrupt the lysosomal pH via interaction with v-ATPase subunits that form a structural pocket within the complex. **(A-C)** Cell lysates (*A, B*) or Lyso fraction (*C*) of 2yr 2N and DS fibroblast were immunoprecipitated (IP) with anti-APP (C1/6.1) antibody, followed by immunoblotting (IB) with antibodies of v-ATPase V0 subunits (*A, C*) and V1 subunits (*B, C*). The values at the bottom of the left IP blot indicate the relative intensities of IP-ed V0a1 and V0d1 normalized by IP-ed APP. The quantification in these experiments is confirmed in two other experiments. **(D)** Cell lysates of 2yr 2N and DS fibroblast were IP with anti-V0a1 antibody, followed by IB with anti-APP (6E10). The values at the bottom of the left IP blot indicate the relative intensities of IP-ed βCTF normalized against IP-ed V0a1. **(E)** *Left panel:* Schematic diagram of *in situ* Proximity Ligation Assay (PLA). PLA performed using V0a1 antibody and C1/6.1 for APP C-terminus in 2yr 2N and DS fibroblasts. The red box represents signal of PLA (Duolink). *Right panel:* Representative fluorescent microscopy image of PLA signals demonstrating association of APP with the V0a1. Scale bar, 10 µm. **(F)** The graph show number (#) of dots per cell (n ≥ 173 cells, three independent). **(G)** Cell lysates of 2yr 2N and DS fibroblast were incubated with varying amounts of each peptide (Pep-1, Pep-2, and Pep-3) for 24 hr, then IP with anti-APP (C1/6.1) antibody, followed by IB with anti-V0a1 antibody. The values at the bottom of the left IP blot indicate the relative intensities of IP-ed V0a1normalized against IP-ed CTFs. **(H)** Lysosomal pH of 2yr 2N and DS fibroblasts treated with either DMSO (-) or 5 µM peptides for 24 hr determined by LysoSensor Y/B dextran (n=3, three independent, triplicate). **(I)** Cell lysates of 2yr 2N and DS fibroblast were IP with anti-pY^682^ APP antibody, followed by immunoblotting with antibody of V0a1. Non-specific Immunoglobulin G (IgG) used as a control IP against 2N cells. The values at the bottom of the left IP blot indicate the relative intensities of IP-ed V0a1 normalized against IP-ed pY^682^βCTF. The quantification in this experiment is confirmed in two other experiments. Actin and GAPDH served as a loading control and LAMP2 served as a lysosomal marker. **(J)** *Left panel:* Schematic diagram of binding with V0a1 and pY^682^APP-βCTF. *Right panel:* The surface of the pYENPTY binding pocket of V0a1. The positions of phosphorylated pYENPTY motif (pY^682^) and F196 and R167 of V0a1 subunits are labelled. **(K)** The binding poses of the pYENPTY motif in the pocket of V0a1. Hydrogen bond and salt bridge interactions by S151, S153, R198, and T250 are labeled with yellow dash lines. The methylene units of R167 and F170 form 1[- hydrophobic contacts with the aromatic ring of the phosphorylated tyrosine. Proline at the pY+3 position has 1 [- hydrophobic interaction with F196. The pYENPTY motif is yellow, and the unsolved region is light blue. Quantitative data (F and H) are presented as mean values with ±SEM, two-tailed unpaired t test (F), ordinary one-way ANOVA with Šidák’s multiple comparisons test (H); ***, P < 0.0005; ****, P < 0.0001. Each dot represents # of dot per field (F) or average value of each independent experiment (H).

Next, to test whether the V0 subunit interacts with flAPP or βCTF, we performed co-IP assays using V0a1 antibody. V0a1 pulled down APP-βCTF, but not flAPP (Fig. 4D). In addition, we confirmed the increased interaction between APP-βCTF and V0a1 in DS fibroblasts compared with 2N cells by an *in situ* proximity ligation assay (PLA) (*34*) using a Duolink technology involving two primary antibodies to V0a1 and APP (C1/6.1) followed by the addition of secondary antibodies fused to oligonucleotide proximity probes (Fig. 4E, left diagram). The proximity probes ligate with connector oligonucleotides to form a circular DNA strand when two proteins are within 40 nm of each other, allowing signal amplification and subsequent binding of fluorescently labeled probes. PLA fluorescence (red dots) was detected in DS fibroblasts at considerably higher levels than in controls (Fig. 4, E to F).

The C-terminal region of APP contains the ^682^YENPTY internalization motif with an NPXpY element, a typical internalization signal for membrane proteins (*35*). The conserved motif is important for interactions with specific cytosolic proteins, including Fe65-PTB2 and Grb2-SH2 that regulate APP metabolism and signaling (*36, 37*). Among three Tyr residues present in the AICD, only Tyr^682^ is phosphorylated *in vivo* (*36*). Both APP phosphorylation and total βCTF levels are increased in AD brain (*38, 39*). A rise in total βCTF will elevate cell levels of pY^682^APP-βCTF, irrespective of whether APP phosphorylation state increases above its constitutive level. To investigate the interaction between V0a1 and the ^682^YENPTY motif, we synthesized an unphosphorylated peptide (QNGYENPTY, Pep-1), a phosphorylated peptide (QNG^P^YENPTY, Pep-2), and a control peptide (QNG^P^YENVTY, Pep-3). Pep-3 is expected to reduce binding to partner proteins due to a conformation change caused by the loss of β-turn conformation at pY+3 (*36*). Cell lysates from 2N and DS fibroblasts were incubated with each peptide for 24 hr at 4°C and then IP-ed with C1/6.1 antibody. The increased APP binding with V0a1 in DS fibroblasts was decreased by only Pep-2 and this occurred in a dose dependent manner (Fig. 4G). In addition, incubation of 2N fibroblasts for 24h with Pep-2, but not Pep-1, mimicked the effect of βCTF elevation in raising the pH of lysosomes to the abnormally high pH in DS fibroblasts (Fig. 4H). To confirm no effect of Aβ_1-42_ on v-ATPase or lysosomal pH, we performed the same experiments as in Fig. 4G and Fig. 4H by incubating cell lysates (fig. S6A) or intact cells (fig. S6B) respectively with Aβ_1-42_. As expected, Aβ_1-42_ peptide had no effect on binding of APP with V0 subunits (Fig. S6A) or on lysosomal pH (fig. S6B). To confirm specific interaction between phosphorylated APP and v-ATPase subunits, we carried out a series of co-IP analyses in cell lysates using anti-APP (phospho-Tyr^682^; pY682) antibody. The binding of pY^682^CTF with V0a1 subunit was increased in DS fibroblasts (Fig. 4I), as evidenced by the higher level of IP-ed V0a1 subunit.

Based on the foregoing results, we analyzed a known AICD structure complexed with Grb2-SH2 (PDB ID: 3MXC) (*36*) to find the binding region in the V0a1 subunit. The phosphate unit of Tyr^682^ is projected toward a pocket composed of R67, R86, S88, S90, S96, and K109 (fig. S7A). The phosphate group forms a hydrogen bonding interaction with S88, S90, and S96 and salt bridge interaction with R67 and R86. Three methylene units of K109 show Π - hydrophobic contact with the phenyl ring of Y682 (fig. S7A). Thus, the structural features of the phospho-Y (pY) binding pocket provide for electrophilic and hydrophobic interactions and stabilize the pY in the pocket. The V0a1 subunit of mammalian brain v-ATPase (PDB ID: 6VQ7), defined from a cryo-electron microscopy (cryo-EM) structure, was analyzed to locate the potential binding site of the pYENPTY motif (*40*). The cryo-EM structure includes an unsolved region from residue 667 to 712 in the V0a1 subunit (fig. S7B, green), and it possesses three consecutive serine residues (S151, S152, and S153) which are highly conserved among species (fig. S7C). In addition, a basic residue, R198, is located in the middle of the N- and C-terminals of the unsolved region (fig. S7B). Thus, we speculated that this region could form a potential binding pocket for the pYENPTY motif. Moreover, two of three serine residues of the loop could be a part of the binding pocket and provide hydrogen bonding interaction to the phosphate group of the tyrosine residue. Molecular modeling, molecular dynamics (MD) simulation, and docking studies were iteratively conducted to build the pocket and estimate the binding pose of the pYENPTY motif (*41, 42*). The model structure shows that the phosphate group of pY possesses hydrogen bonding interaction with S151, S153, and T250 and has a salt bridge interaction with R198 (Fig. 4, J to K). The phenyl ring of pY forms hydrophobic contacts with the methylene units of R167 and F170 in a way similar to that shown for the Grb2-SH2 and AICD complex (fig. S7A). Furthermore, MD simulation followed by MM-GBSA calculation (*43*) shows the estimated binding affinity of the phosphorylated peptide to the pocket is more stable than its unphosphorylated counterpart (Δ Δ G_bind_ = −85.6 kcal/mol versus −67.8 kcal/mol).

Taken together, these analyses strongly suggest that APP-βCTF interacting specifically with V0a1 modulates docking of the V1 sector to the V0 sector and that Tyr^682^ phosphorylation on the ^682^YENPTY motif of APP plays a crucial role in this interaction.

### Decreasing the levels of phospho-Y^682^ APP-βCTF rescues v-ATPase dysfunction in DS

In further investigations on the effect of Tyr^682^ phosphorylation on v-ATPase function, we found, as expected, elevated levels of pY^682^ APP-βCTF in DS fibroblasts (Fig. 5A). Given the possibility that Fyn kinase, which binds the ^682^YENPTY motif of APP and increases APP tyrosine phosphorylation in AD neurons (*44*), represents *one possible source* for the elevation of pY^682^ APP-βCTF levels in DS fibroblasts, we tested whether Saracatinib (AZD0530), a Src family of nonreceptor tyrosine kinases inhibitor with high potency for Fyn, would lower pY^682^APP-βCTF levels and, most importantly, rescue v-ATPase deficits. Testing with various concentrations of AZD0530 revealed a partial reduction of phospho-Y^420^-Fyn (pFyn) at 2 µM AZD0530 (fig. S8A) indicating inhibited activity. Next, measurements of lysosomal pH and v-ATPase activity in 2N and DS fibroblasts after treating with or without AZD0530 demonstrated rescue of elevated lysosomal pH, v-ATPase activity deficits, and reduced lysosomal targeting of V1D (Fig. 5, B to E) in AZD0530 treated DS fibroblasts, which also exhibited lowered pY^682^APP-βCTF (Fig. 5A). In addition, decreased interaction between V0a1 and APP-βCTF was also demonstrated in DS fibroblasts treated with AZD0530 (Fig. 5F).

**Fig. 5.**
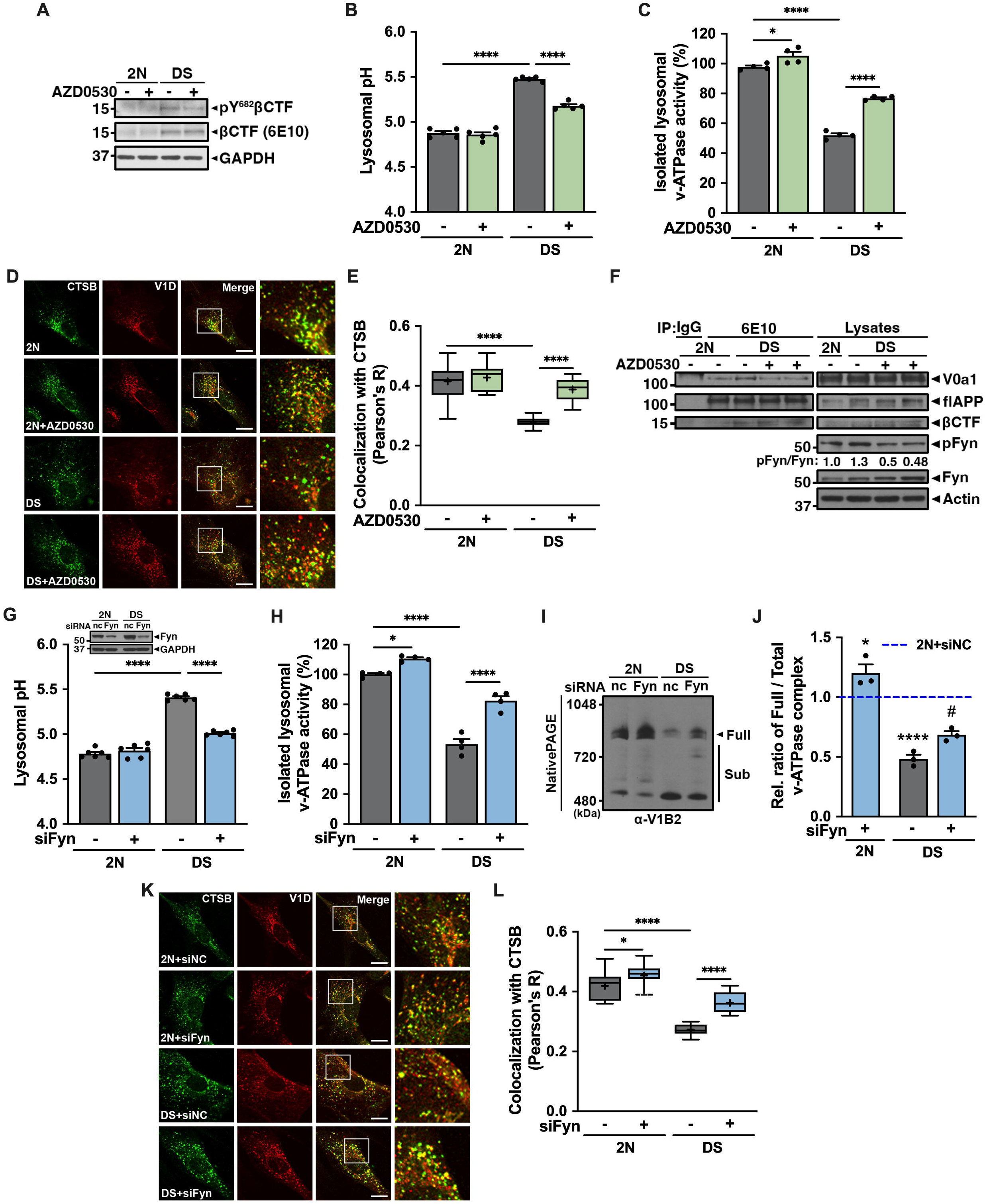
Decreased levels of phosphorylated Tyr^682^ APP-βCTF rescues v-ATPase dysfunction in DS fibroblasts. **(A)** Immunoblots of phospho-Y^682^βCTF and βCTFs distribution in total lysates of 2yr 2N and DS fibroblasts. **(B)** Lysosomal pH of 2yr 2N and DS fibroblasts treated with either DMSO (-) or 2 µM a Src family of nonreceptor tyrosine kinase inhibitor (AZD0530) for 24 hr determined by LysoSensor Y/B dextran (n=5, five independent, triplicate). **(C)** ATP hydrolysis activity of v-ATPase measured using lysosomal fractions from DMSO or AZD0530 treated 2yr 2N and DS fibroblast (n=4, four independent, triplicate). **(D, E)** Double-immunofluorescence labeling shows colocalization of V1D and CTSB in 2yr 2N and DS fibroblasts treated with DMSO or AZD0530 (*D*). Scale bar, 10 µm. Quantification of v-ATPase V1D and lysosomal marker, CTSB, shows colocalization in 2N and DS fibroblasts treated with DMSO or AZD0530 as calculated by Pearson’s correlation coefficient (n ≥ 118 cells, three independent) (*E*). **(F)** After treated with DMSO (-) or 2 µM AZD0530 (+), cell lysates of 2yr 2N and DS fibroblast were IP with anti-APP (6E10) antibody, followed by IB with anti-V0a1 antibody. **(G)** Lysosomal pH of 2yr 2N and DS fibroblasts transfected with siRNA for either negative control (-) or 100 nM siFyn (+) for 48 hr, as determined by LysoSensor Y/B dextran (n=6, six independent, quadruplicate). Immunoblot represents Fyn kinase protein levels in 2N and DS fibroblast. Actin or GAPDH served as a loading control. **(H)** ATP hydrolysis activity of v-ATPase measured using lysosomal fractions from siRNA transfected 2yr 2N and DS fibroblast (n=4, four independent, triplicate). **(I)** Membrane fractions from siRNA transfected 2yr 2N and DS fibroblasts were resolved using the native PAGE. **(J)** The graph represents relative ratio of full complex divided by total (full plus partial complexes) complexes of v-ATPase. **(K, L)** Double-immunofluorescence labeling shows colocalization of V1D and CTSB in siRNA transfected 2yr 2N and DS fibroblasts (*K*). Scale bar, 10 µm. Quantification of v-ATPase V1D and lysosomal marker, CTSB, shows colocalization in siRNA transfected 2N and DS fibroblasts, as calculated by Pearson’s correlation coefficient (n ≥ 139 cells, three independent) (*L*). Quantitative data (B, C, E, G, H, J, and L) are presented as mean values with ±SEM, ordinary one-way ANOVA with Šidák’s multiple comparisons test; *, P < 0.05; ****, P < 0.0001. Statistically significance between groups is shown by symbols: *2N vs others, #DS vs DS+AZD0530/siFyn. Each dot represents average value of technical replicates from each independent experiment.

To confirm the effects of lowering pY^682^APP-βCTF by AZD0530 on the assembly of v-ATPase, small interference RNA constructs were used to knock down the expression of Fyn (fig. S8B). As expected, siFyn, but not siNC, substantially restored lysosomal acidification (Fig. 5G), v-ATPase activity (Fig. 5H), and lysosomal targeting of V1D (Fig. 5, K to L) in DS fibroblasts. Fyn knockdown also induced reassembly of disrupted V0/V1 complexes in DS fibroblasts (Fig. 5, I to J). Despite higher v-ATPase activity (Fig. 5H) and assembly (Fig. 5, I to J) in 2N cells, lysosomal pH was not lowered to an abnormal acidic range (Fig. 5G) consistent with the expectation that complementary regulatory systems, such as CLC-7 and additional proton leak channels (*14*), compensate for acute proton import changes to maintain optimal lysosomal pH. A lowering of pY^682^APP-βCTF level below normal baseline would likely come into play not to lower pH below the normal range but to correct minor rises in lysosomal pH under physiological stress (or mild disease) conditions.

Collectively, these data indicated that the dysfunction of v-ATPase in DS fibroblasts is likely mediated by elevated levels of pY^682^APP-βCTF and that reducing its levels restores v-ATPase functioning.

### v-ATPase dysfunction in DS model mice and human DS brain reflect impaired association of V0 and V1 sectors

We next extended our observations *in vivo* by investigating the possibility of similar lysosomal v-ATPase dysfunction and defective acidification in the Ts[Rb(12.17^16^)]2Cje (Ts2) mouse model of DS (*45*). Villar et al., identified and characterized Ts2 mice, carrying a chromosomal rearrangement of the Ts65Dn genome. The Ts2 mouse expresses the same complement of trisomic genes as Ts65Dn and displays the same overt DS-related phenotype (*5*), but has the advantages of yielding male mice that are fertile (unlike Ts65Dn) and female mice that have higher trisomy transmission rates, resulting in a ∼3-fold higher viable offspring rate compared to that in Ts65Dn mice (*45*). Extensive published analyses have demonstrated that Ts2 exhibits an identical neurological phenotype to Ts65Dn, with indistinguishable AD-related pathological and synaptic plasticity abnormalities, changes in abnormal APP metabolism, age-dependent endosomal and cholinergic phenotypes, and timings of onset and progression of these anomalies (*46-48*).

To monitor lysosomal acidification *in vivo*, we crossed Ts2 mice with Thy1 promotor-driven transgenic mRFP eGFP LC3 (TRGL6) mice to yield (Ts2*TRGL) mice as described previously (*13*). Introduction of a third fluorescence probe by immunocytochemical labeling with cathepsin D (CTSD) antibody allowed us to detect lysosomes and autolysosomes (AL), and, based on their acidification state, to discriminate LC3-positive AL that had fused with lysosomes but had not properly acidified from LC3-positive, cathepsin D-negative autophagosomes. A computer assisted unbiased measurement of hue angle (an index of wavelength) of individual triple fluorescent vesicles enabled a relative vesicle pH determination and objective assignment of vesicle identity (*13*). The confocal images were collected from neocortex sections of 8-9 months old Ts2*TRGL and age-matched TRGL mice (Fig. 6A). In Ts2*TRGL mice, numbers of fully acidified AL (purple puncta) were decreased compared to that in control TRGL mice, whereas poorly acidified AL (white puncta) increased in number (Fig. 6B).

**Fig. 6.**
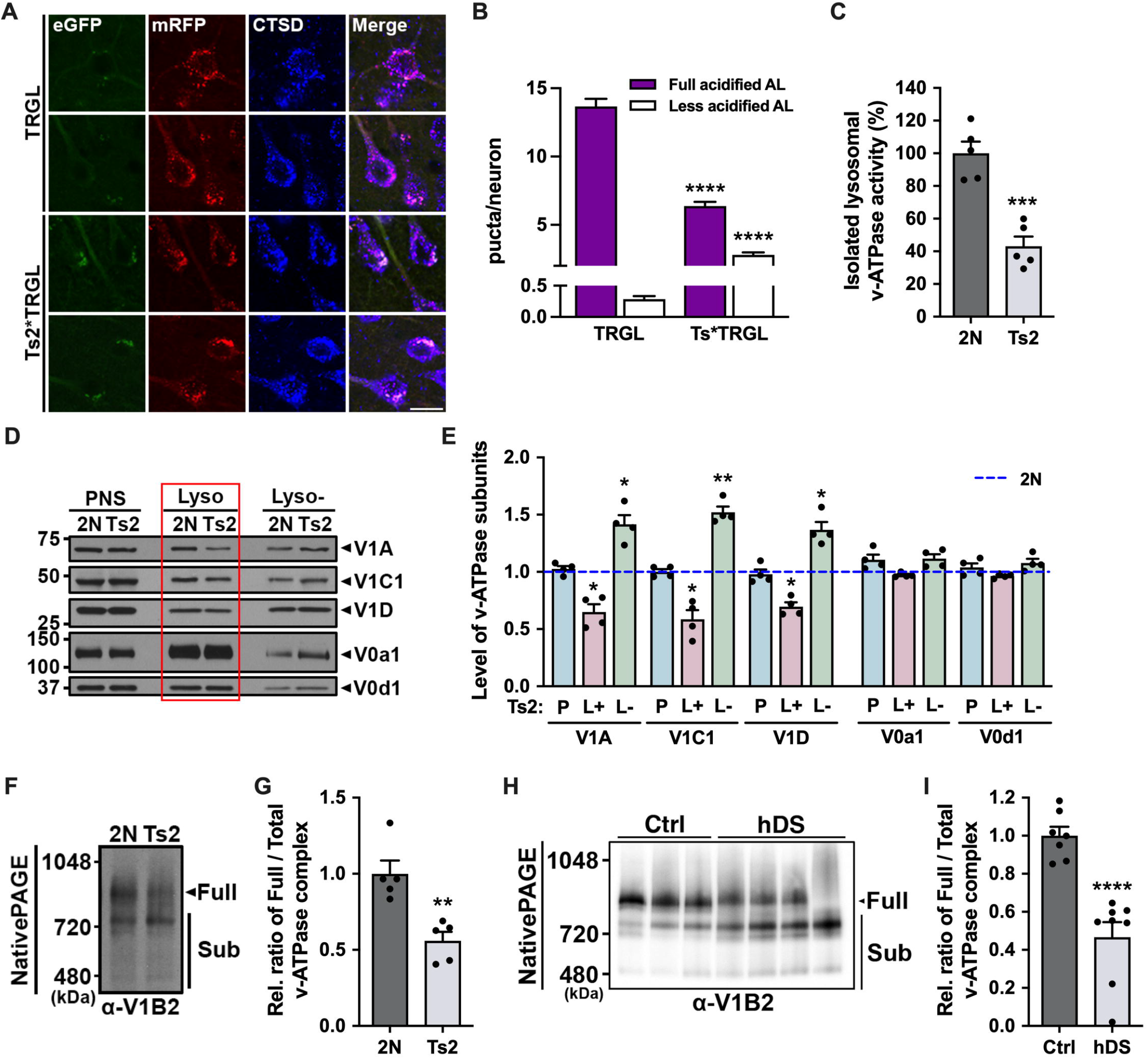
Defective lysosomal v-ATPase function in Ts2 mouse model and human DS brain. **(A, B)** Representative fluorescence images of tfLC3 color changes of TRGL or Ts2*TRGL adult brain (*A*) and respective quantification of puncta representing autolysosomes (fully or less acidified) (*B*). For data analysis, color change of tfLC3-positive vesicles was calculated with CTSD co-labeling. Bar colors denote the colors of puncta; purple bars indicate tfLC3 with CTSD (fully acidified ALs, quenched eGFP) and white bars indicate tfLC3 with CTSD (less acidified ALs, unquenched eGFP) (n=504 neurons, 6 TRGL mice; n=537 neurons, 6 Ts2*TRGL mice). **(C)** ATP hydrolysis activity of v-ATPase was measured using Lyso fraction (15-18) of the iodixanol step gradient from adult mice (n=5, triplicate). **(D)** Adult mouse brain from control (2N) and Ts2 homogenates were fractionated through an iodixanol step gradient. Each fraction (lysosomal enriched fraction [Lyso]: 15-18; exclude lysosomal enriched fraction [Lyso-]: 1-14 & 19-22) and post-nuclear supernatant (PNS) were resolved using the SDS-PAGE and immunoblotted with anti-V1A, V1C1, V1D, V0a1, and V0d1 antibodies. **(E)** The graphs represent levels of v-ATPase subunits from each fraction (n=4). **(F, H)** Membrane fractions from mouse brain (*F*) or frozen human brain tissue (*H*) were resolved using the native. **(G, I)** The graph represents relative ratio of full complex divided by total (full plus partial complexes) complexes of v-ATPase (n=5). Quantitative data (B, C, E, G, and I) are presented as mean values with ±SEM, two-tailed unpaired t test (C, G, and I), ordinary one-way ANOVA with Šidák’s multiple comparisons test (B and E); *, P < 0.05; **, P < 0.005; ***, P < 0.0005; ****, P < 0.0001. Each dot corresponds to average value of technical replicates from one mouse.

Next, to assess lysosomal v-ATPase activity in these mice, we performed biochemical fractionations to obtain Lyso fractions using an iodixanol step gradient (fig. S9). WB analysis using various organelle markers identified Lyso within fractions 15-18 (fig. S9; Lyso), while measurements of the ATP-hydrolytic activity in this enriched lysosome fraction revealed a ∼50% lowered ATP hydrolysis rate in the lysosomes from Ts2 mice relative to those in 2N control mice (Fig. 6C). While the TRGL probe assessed pH selectively in neurons, the v-ATPase assay assessed activity from all brain cell types, which are expected also to have a Ts2 phenotype in this systemic disorder.

We also performed Western blot analysis using iodixanol step gradient samples to investigate whether lysosomal targeting of v-ATPase subunits to lysosomes was also disrupted in Ts2 mice. V1 subunits in the Lyso (L+) fraction from Ts2 mouse brain were present at markedly decreased levels compared to those in 2N and were increased in other fractions (Lyso-; L-) (Fig. 6, D to E). V0 subunit levels in the Lyso fraction were not changed (Fig. 6, D to E), as expected from our experiments above. To confirm disruption of v-ATPase assembly in Ts2 mouse, we performed native PAGE analysis using a membrane fraction. Native PAGE analysis showed that the relative ratio of fully assembled v-ATPase divided by total v-ATPase (fully assembled v-ATPase plus subcomplexes of v-ATPase) was significantly reduced in the Ts2 mouse, compared with 2N (Fig. 6, F to G). We additionally observed disruption of v-ATPase assembly in the membrane fraction from frozen brain from individuals with DS, which showed increased levels of flAPP and βCTF (fig. S10), compared to the control group (Fig. 6, H to I).

### Lysosomal dysfunction in DS model mice is reversed by lowering elevated pY^682^ APP-βCTF levels

In further investigations, we showed that the level of pY^682^APP-βCTF, measured with Tyr phosphorylation specific antibodies, is nearly 2-fold higher in Ts2 mouse brain by WB analysis (Fig. 7, A to B). To determine the effects of raised levels of pY^682^APP-βCTF, *in vivo*, we then treated Ts2 mice with vehicle or 5 mg/kg/day AZD0530 by oral gavage for 4 weeks (*49*). We confirmed by WB analysis that AZD0530 decreased pY^682^APP-βCTF levels in Ts2 mice *in vivo* (Fig. 7, C to D) and significantly rescued abnormal lysosomal acidification (Fig. 7, E to F) compared to vehicle treated (Veh) mice and restored v-ATPase activity (Fig. 7G). In addition, the lowered level of pY^682^APP-βCTF in Ts2 mice restored the normal ratio of full versus total v-ATPase complex in Ts2 mouse brain (Fig. 7, H to I) and increased association of V1 subunits in Lyso fractions (Fig. 7, J to K). Notably, supporting our foregoing evidence in fibroblasts for tonic modulation of v-ATPase, the lowered pY^682^APP-βCTF levels in wild-type mice induced supra-normal levels of v-ATPase activity and proportions of fully assembled v-ATPase complex in enriched lysosome fractions and an increased proportion of V1 subunits associated with the V0 sector on lysosomes.

**Fig. 7.**
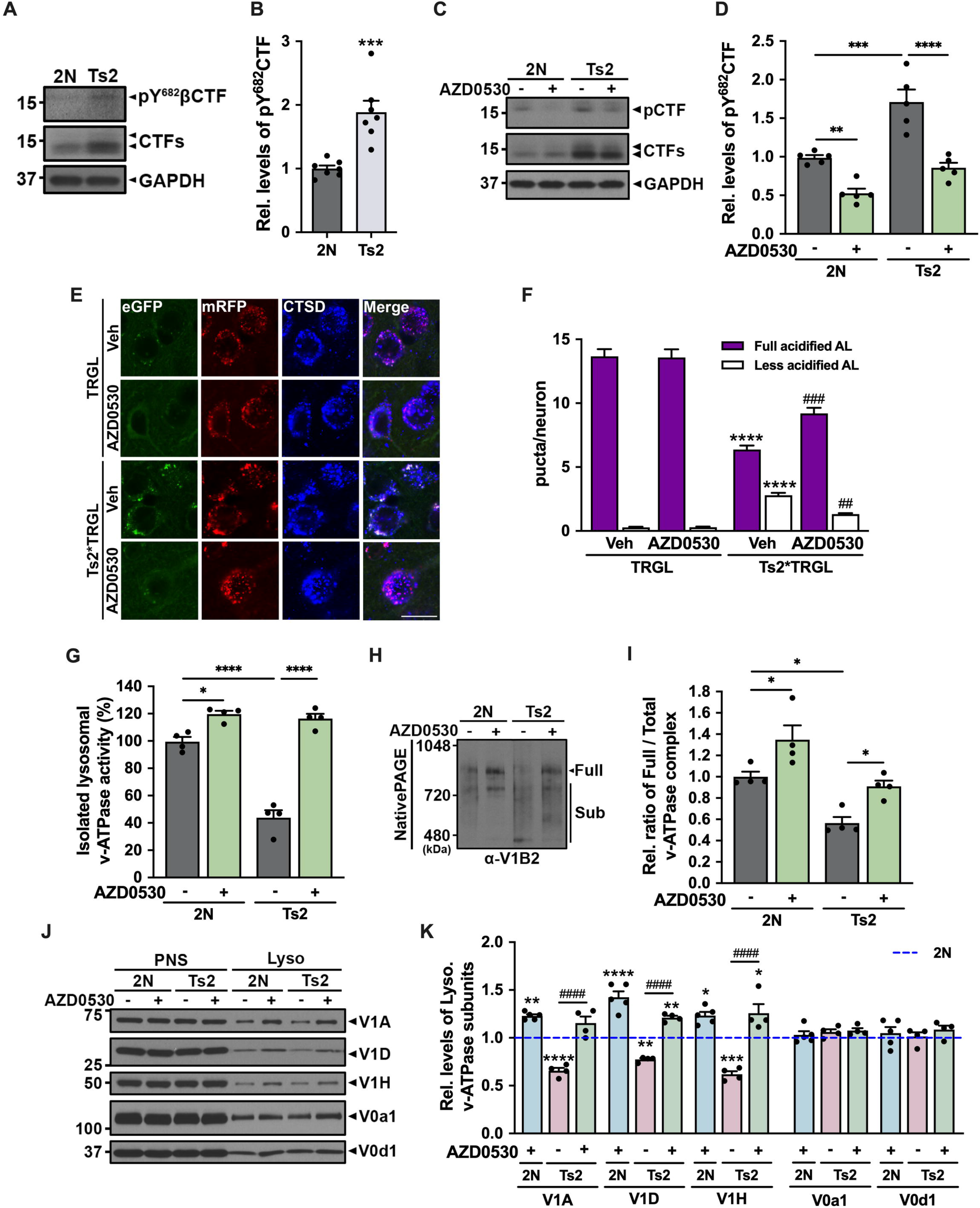
Reduction in levels of phosphorylated Tyr^682^ APP-βCTF *in vivo* reverses v-ATPase dysfunction in brains of DS model mice. **(A)** Immunoblot represents phospho-Y^682^CTF and CTFs levels in 2N and Ts2 mouse brain. **(B)** The graphs show band intensity of pY^682^βCTF protein (n=7). **(C)** The immunoblot represents phospho-Y^682^CTF (pCTF) and CTFs levels in 2N and Ts2 after treated with either vehicle (-) or AZD0530 (+) for 4 weeks. GAPDH served as a loading control. **(D)** The graphs show band intensity of phospho-Y^682^CTF (n=5). **(E, F)** Representative fluorescence images of tfLC3 color changes of adult brain after treatment with vehicle (Veh) or AZD0530 for 3 weeks (*E*) and respective quantification of puncta representing autolysosomes (fully or less acidified) (n=452 neurons, 4 TRGL mice; n=491 neurons, 4 AZD0530 treated TRGL mice; n=476 neurons, 4 Ts2*TRGL mice; n=484 neurons 4 AZD0530 treated Ts2*TRGL mice) (*F*). **(G)** ATP hydrolysis activity of v-ATPase measured using Lyso fraction (15-18) of iodixanol step gradient from adult mouse brain after treatment with vehicle (-) or AZD0530 (+) for 4 weeks (n=4). **(H)** Membrane fractions from AZD0530 treated adult mouse brain were resolved using native PAGE and immunoblotted with anti-V1B2 antibody. **(I)** The graph represents relative ratio of full complex divided by total (full plus partial complexes) complexes of v-ATPase (n=4). **(J)** Homogenates from AZD0530 treated 2N and Ts2 adult mice brain were fractionated through an iodixanol step gradient. The Lyso fraction and the PNS were resolved using the SDS-PAGE and immunoblotted using indicated antibodies. **(K)** The graphs represent levels of lysosomal v-ATPase subunits (n=5 TRGL mice; n=5 AZD0530 treated TRGL mice; n=4 Ts2*TRGL mice; n= 4 AZD0530 treated Ts2*TRGL mice). Quantitative data (B, D, F, G, I, and K) are presented as mean values with ±SEM, two-tailed unpaired t test (B), ordinary one-way ANOVA with Šidák’s multiple comparisons test (D, F, G, I, and K); *, P < 0.05; **, P < 0.005; ***, P < 0.0005; ****, P < 0.0001. Statistically significance between groups is shown by symbols: *2N+Veh vs others, #DS+Veh vs DS+AZD0530. Each dot corresponds to average value of technical replicates from one mouse.

The foregoing results establish lysosomal v-ATPase dysfunction in DS model brain *in vivo* as well as in authentic DS in fibroblasts from individuals with DS. We also demonstrate its further dependence on Tyr^682^ phosphorylation of APP in DS.

### v-ATPase deficits are associated with pY^682^APP-βCTF in APP-based mouse models of AD

As an initial extension of our findings in DS to early onset AD models induced by *hAPP* over-expression or *APP/PSEN1* mutations, we investigated the relevance of Tyr^682^APP phosphorylation to defective lysosomal acidification and function in two AD mouse models: 6-month-old 5xFAD mice exhibiting early-onset amyloidosis (at 2 months) and 6-month-old Tg2576 mice with a later age of onset of amyloidosis (at 10-12 months). v-ATPase activity measured in Lyso fractions was reduced by ∼45% in each model relative to WT controls (Fig. 8A and 8F). Phosphorylation of APP at Tyr^682^ is increased in AD brain (*38, 39, 44*). Accordingly, we found levels of pY^682^ APP-βCTF to average nearly 3-fold higher in Tg2576 mouse brain (Fig. 8, B to C). To confirm disruption of v-ATPase assembly in Tg2576 mouse, we performed native PAGE analysis using membrane fractions from cortex. Native PAGE analysis showed that the relative ratio of fully assembled v-ATPase divided by total v-ATPase complex was significantly reduced in the Tg2576 mouse, compared with WT (Fig. 8, D to E). We observed similar results in brains of the 5xFAD mouse model of AD (Fig. 8, F to J).

**Fig. 8.**
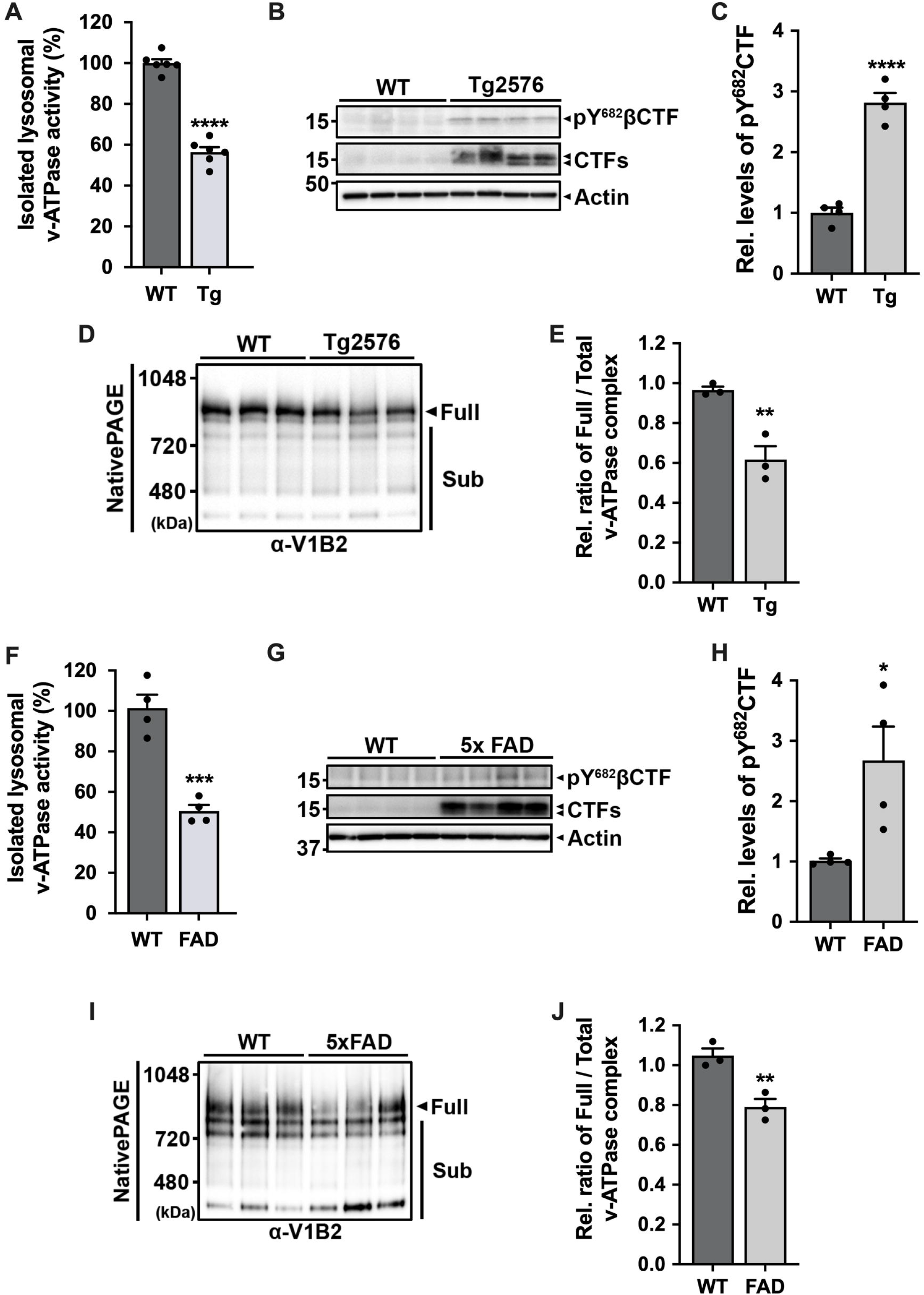
Defective v-ATPase dysfunction in APP-based AD model mouse. **(A)** Lysosomal v-ATPase activity measured colorimetrically as ATP hydrolysis with and without ConA. Activity assays were performed on lysosomal enriched fraction (15-18) of iodixanol step gradient from 6-months Tg2576 female adult brain pre-treated with *o*-vanadate to minimize nonspecific ATPase activity (n=6, triplicate). **(B)** Immunoblot represents pY^682^CTFs and APP-CTFs protein levels in 6-months female Tg2576. Actin served as a loading control. **(C)** The graph shows relative band intensity of pY^682^CTF proteins (n=4). **(D)** Membrane fractions from 6-months Tg2576 female adult brain were resolved using the native PAGE and immunoblotted with anti-V1B2 antibody. (E) The graph represents relative ratio of full complex divided by total (full plus partial complexes) complexes of v-ATPase (n=3). **(F)** Lysosomal v-ATPase activity measured colorimetrically as ATP hydrolysis with and without ConA. Activity assays were performed on lysosomal enriched fraction (15-18) of iodixanol step gradient from 6-months 5xFAD female adult brain pre-treated with *o*-vanadate to minimize nonspecific ATPase activity (n=4, triplicate). **(G)** Immunoblot represents pY^682^CTFs and APP-CTFs protein levels in 6-months female 5xFAD. **(H)** The graphs show band intensity of pY^682^CTF protein (n=4). **(I)** Membrane fractions gradient from 6-months 5xFAD female adult brain were resolved using the native PAGE and immunoblotted with anti-V1B2 antibody. **(J)** The graph represents relative ratio of full complex divided by total (full plus partial complexes) complexes of v-ATPase (n=3). Quantitative data (A, C, E, F, H, and J) are presented as mean values with ±SEM, two-tailed unpaired t test; *, P < 0.05; **, P < 0.005; ***, P < 0.0005; ****, P < 0.0001. Each dot corresponds to average value of technical replicates from one mouse.

## Discussion

In all forms of AD, the impaired clearance of autophagic and endocytic substrates by lysosomes causes profuse accumulations of neuronal waste, including the build-up in lysosomes of key APP metabolites (eg. βCTF, Aβ), ultimately leading to lysosomal membrane permeability and neuronal cell death (*26, 50-52*). In this report, we have established a primary APP-dependent mechanism underlying lysosome dysfunction in DS, a congenital genetic disorder responsible for the most prevalent form of early onset AD (*53-55*). Lysosomal v-ATPase function is disrupted by APP-βCTF directly interacting with the critical membrane-anchored V0a1 subunit of v-ATPase, thereby impeding assembly of the complete V1 sector with the V0 sector of v-ATPase. The relevance of this molecular interaction is supported by a previous unbiased analysis of the synaptosome interactome of the APP intracellular domain (AICD) harboring the conserved YENPTY region of interest. This analysis revealed V0a1 and V0d1 and various V1 subunits to be AICD interactome members (*33*). Reinforcing the pathogenic importance of the APP-βCTF interaction with V0a1 subunit, mutations of *PSEN1*, a major cause of early-onset AD (*56*), are also known to disrupt PSEN1 holoprotein ER chaperone role in facilitating V0a1 glycosylation, folding, and stability, thereby impeding adequate delivery of the key subunit to lysosomes (*25, 57*). Further highlighting V0a1 as a common disease target in AD, other evidence suggests that V0a1 subunit dysfunction is compromised in additional neurodegenerative conditions featuring prominent lysosomal dysfunction, including LRRK2 linked PD (*58*) and the lysosomal storage disorder, CLN1 (*59*).

We further establish here that Tyr^682^ APP phosphorylation confers selectivity to the pathogenic interaction of pY^682^APP-βCTF with V0a1. The modestly elevated level of endogenous APP-βCTF in DS (*60*) is necessary and *sufficient* to raise lysosomal pH and partially inactivate lysosomal enzymes, as these pathological consequences are reversed by reducing APP-βCTF levels (*5*) either directly or via BACE1 inhibition, which also lowers pY^682^APP-βCTF levels. Alternatively, reversal could also be achieved by inhibiting the activity of tyrosine kinases, which targets Tyr^682^ on APP (*61-63*) and is over-active in DS. While other genes in the trisomic chromosomal region in DS fibroblasts and Ts2 mice may have additional influence on lysosomal function, we show that elevated levels of pY^682^APP-βCTF in AD mouse models, which lack the trisomy of DS, induce the same lysosomal acidification deficit as in DS models.

We found inhibition of Fyn activation to be a contributing factor in lowering pY^682^ APP-βCTF levels in DS, consistent with other studies in DS and AD (*39, 64*). We do not contend, however, that Fyn is the only possibly relevant kinase involved in all lysosomal pH-related disease contexts, or even that *over-activation* of tyrosine kinase is essential to the mechanism of v-ATPase deficiency operating in AD. Tyr^682^ APP phosphorylation is a constitutive process, so a proportion of APP and βCTF molecules normally contains Tyr^682^ phosphate, as we have shown. Levels of pY^682^ APP-βCTF may therefore become elevated without over-activating any particular tyrosine kinase if (a) β-cleavage of APP is accelerated (eg. many AD-related factors that activate BACE-1); (b) conversion of βCTF to Aβ is inhibited (eg. PSEN1/2 loss of function or inhibition); or (c) βCTF is induced to accumulate in lysosomes (eg. cathepsin inhibition in various disease contexts). In our study, we used an inhibitor with high affinity for Fyn kinase as a tool to suppress Tyr^682^ phosphorylation in order to provide proof of principle that lowering levels of pY^682^ APP-βCTF, regardless of the specific cause of an abnormal elevation, can ameliorate v-ATPase dysfunction.

The pathogenic binding to v-ATPase is an effect of APP-βCTF, but not Aβ or APP-⍰CTF, based on our use of γ- and ⍰-secretase inhibitors and peptide competition assays with v-ATPase subunits. Brain levels of APP-βCTF are not only elevated in DS but also in individuals with familial Alzheimer’s Disease (FAD) caused by APP or PSEN1 mutations (*5, 29, 65, 66*) and in IPSCs derived from homozygous APOE4 carriers (*67*). Importantly, steady-state levels of APP-βCTF likely underestimate its production and flux through the endosomal-lysosomal system when APP endocytosis rates are elevated, as is likely in AD and DS. In DS, expression of the AD phenotype of endosomal and lysosomal anomalies (*5, 29, 66*) is dependent on a single extra copy of APP, highlighting the sensitivity of the v-ATPase system response to modest APP-βCTF elevations. Levels of pY^682^ APP-βCTF in brain are also elevated in individuals with late-onset “sporadic” AD (*38, 39*) as well as in two different AD model mice (Tg2576 and 5xFAD).

The foregoing mechanism of v-ATPase impairment in FAD/DS is potentially relevant to explaining the similar lysosomal dysfunction in sporadic AD. In late-onset (sporadic) AD, APP-βCTF elevation, in the absence of a rise in APP levels, results from potentially multiple factors that may be additive. These include an increased level of BACE1 activity (*68, 69*), cell stress-activated Tyr^682^ APP phosphorylation, which notably also promotes β-cleavage of APP over alpha-cleavage (*70*), and APP processing regulation by GWAS identified AD risk genes such as BIN1, SORL1, ABCA7, and RIN3 (*71, 72*). Additionally, as hydrolytic efficiency declines in aging due to rising oxidative stress, lipid accumulations, and other aging factors (*73-75*), APP-βCTF may well build-up in lysosomes disproportionately to flAPP levels (*76, 77*) and inhibit v-ATPase.

Besides regulating substrate hydrolysis, the intraluminal pH of lysosomes modulates diverse signaling functions including maturation of lysosomal hydrolases (*78*), modification of certain cargoes delivery to lysosome (*79, 80*), fusion of lysosomes with other organelles/cell surface (*81, 82*), and retrograde axonal transport of LAMP-positive compartments (*83*). Therefore, v-ATPase activity is increasing recognized to be tightly regulated (*21, 84*). Varied factors modulate V1 sector association with the V0 sector (*24*) and reversible association of specific individual V1 subunits with the complex may also have modulatory effects on activity (*17, 85*). The regulatory factors are best characterized in yeast (*17, 18, 84*) although several factors have been reported in mammals (*86-88*). Given the diverse biological roles of v-ATPase and luminal pH regulation, it is not surprising to find global effects of loss of function mutations of v-ATPase in various diseases, especially those with prominent neurodegenerative phenotypes (*24*). The association of early lysosome acidification failure in AD models with a cascade of autophagic and neurodegenerative phenomena underlying β-amyloidosis and lysosomal cell death has been recently described (*26*).

Although not the main focus of our analyses on CTF pathogenicity in AD, we observed consistently that manipulations lowering pY^682^APP-βCTF below normal baseline, induced supra-normal v-ATPase assembly, lysosomal ATPase activity, and acidification. Such conditions included knockdown of APP (Fig. 2), BACE1 inhibitor treatment (Fig. 3), and reducing levels of tyrosine kinase activity (Fig. 5 and 7). Collectively, these findings support the possibility that change in APP-βCTF levels in normal lysosomes under varying physiological levels may tonically modulate v-ATPase activity, although implications of such potential modulation, including therapeutic applications, require future investigation.

In conclusion, *APP* joins *PSEN1* as not only the major monogenic causes of early-onset AD but as genes directly responsible for extreme disruption of lysosomal function and autophagy that is evident in essentially all forms of AD (*89*) and shown to be the catalyst and driver for varied downstream pathophysiology associated with AD development (*26, 90, 91*). Our results unequivocally establish the importance of APP-βCTF in AD etiology by demonstrating a direct pathogenic action of APP-βCTF in the context of DS and AD. Together with other emerging data, our findings point toward acidification failure within the lysosomal network as a potential unifying pathogenic mechanism in AD, driven by converging genetic, aging, and environmental factors of varying influence depending on the subtype of AD (*24, 73, 75*). Finally, mounting preclinical evidence establishes the ameliorative impact of lysosome re-acidification not only on lysosomal efficacy but on neuropathological signatures, synaptic plasticity, and cognition. As such, acidification rescue represents a promising innovative target for the remediation of lysosome dysfunction in AD and related disorders (*92-94*).

## Materials and Methods

### Materials

Materials are provided in Supplementary Table S1.

### Cell lines and transfection

Human foreskin fibroblasts from Down syndrome patient (DS) and diploid age-matched controls (2N) were purchased from the Coriell Cell Repositories (AG06922, AG07095, AG07096, GM08680) and maintained according to the distributor’s protocols (https://www.coriell.org/). Throughout this study, fibroblasts were used below passage number 15 to keep the original character and morphology.

siRNA against human APP and negative control DsiRNA were purchased from Integrated DNA Technologies (IDT) and used as previously described (*31*). siRNA against human Fyn was purchased from Life Technologies Corp. Cells were transfected using Lipofectamine RNAiMAX according to the manufacturer’s protocol.

### Animals

All animal procedures were performed following the National Institutes of Health Guidelines for the Humane Treatment of Animals, with approval from the Institutional Animal Care and Use Committee at the Nathan Kline Institute for Psychiatric Research. Four mouse models were used: (i) Control mice (2N), C57BL/6JEi x C3H/HeSnJ; (ii) Ts[Rb(12.17^16^)]2Cje (Ts2), carrying a chromosomal rearrangement of the Ts65Dn genome whereby the marker chromosome has been translocated to Chromosome 12 (MMU12) forming a Robertsonian chromosome (*45*), were studied at 8-9 months; (iii) TRGL6 (Thy1 mRFP-eGFP-LC3, Line 6), expressing tandem fluorescent mRFP-eGFP-LC3 under individual Thy 1 promoters (*13*); (iv) Ts2 x TRGL6, a transgenic Ts2 mouse model expressing tandem-fluorescent mRFP-eGFP-LC3 under individual Thy1 promoters, were studied at 8-9 months; (v) 5xFAD x TRGL6, 5xFAD (Tg6799, C57BL/6NTAC), which express mutant human APP and PSEN1 (APP KM670/671NL: Swedish, I716V: Florida, V717I: London, PSEN1 M146L, L286V), were crossed with TRGL6 and were studied at 6 months together with age-matched controls; (vi) Tg2576 (B6;SJF, Tg(APPSWE)2576Kha), expressing mutant human APP (Swedish K670N/M671L), were studied at 6 months together with age-matched controls. All mice were genotyped by PCR. Mice were group-housed under 12 h light/dark cycle at constant room temperature and provided with food and water ad libitum. Age-matched WT mice of comparable background were used as controls.

### Ts2 x TRGL mouse generation

For the generation Ts2 x TRGL6 mice, Ts2 mice were bred with our previously described TRGL6 (***T****hy1* m**R**FP-e**G**FP-**L**C3, Line **6**) transgenic mice expressing tandem fluorescent mRFP-eGFP-LC3 (*13*). Ts2*TRGL mice were studied at 8-9 month together with age-matched controls. Mice used in this study were maintained according to Institutional Animal Care and Use Committee guidelines and protocols for animal experimentation at Nathan S. Kline Institute.

### Analysis of autolysosome acidity of TRGL mouse and Vesicle quantification

Immunohistochemistry was performed as previously described (*25*). Animals were anesthetized with a mixture of ketamine (100 mg/kg body weight) and xylazine (10 mg/kg body weight) and intracardially perfused by 0.9% saline. Brains were dissected and immersed in the same fixative for 24 h and then 40-um sagittal sections were made using a vibratome. Immediately after sectioning, sections were immunolabeled with the indicated antibodies overnight and then visualized with Alexa Fluor conjugated secondary antibodies. Imaging was performed using a plan-Apochromat 40x/1.4 oil objective lens on a LSM880 laser scanning confocal microscope. The 3 colors (RGB; red, green, blue) intensity of each vesicle were calculated by Zen from Carl Zeiss Microscopy using the profile function. The R, G, and B ratio of each vesicle was calculated into a hue angle and saturation range by following formula: Hue° = IF(180/PI()*ATAN2(2*R-G-B,SQRT(3)*(G-B))<0,180/PI()*ATAN2(2*R-G-B,SQRT(3)*(G-B)) +360,180/PI()*ATAN2(2*R-G-B,SQRT (3)* (G-B))). Saturation percent of the hue angle was calculated by entering the values of R, G, and B for a given puncta into the following formula = (MAX(RGB)-MIN(RGB))/SUM(MAX(RGB)+MIN(RGB))*100, provided lightness is less than 1, which is the usual case for our data. Vesicle quantification was performed as previously described (*13*).

### Saracatinib pharmacokinetics

To examine the effect of a Src family of nonreceptor tyrosine kinase inhibitor (Saracatinib; AZD0530) in a preclinical DS mouse model, mice received a dose of 5 mg/kg per day administered orally twice daily for 3-4 weeks

### Lysosomal pH measurement

Following the addition of 250 µg LysoSensor™ Yellow/Blue dextran treatments, cells were incubated for 24 hr. The samples were then read in a Wallac Victor 2 fluorimeter (Perkin Elmer) with excitation at 355 nm. The ratio of emission 440 nm/535 nm was then calculated for each sample. The pH values were determined from the standard curve generated via pH calibration samples. To measure lysosomal pH in presence of peptide, we directly exposed 5 µM peptide to the cells for 24 hr at 37 ºC, wash out the peptide containing medium (three times), and then added 250 µg LysoSensor™ Yellow/Blue dextran for 24 hr at 37 ºC.

### Lysosomal isolation

Cells were incubated in growth medium containing 10 % Dextran conjugated magnetite (DexoMAG™ 40) for 24 hr, then chased in normal growth media for 24 hr. Cells were washed with 1X PBS then harvested in 4 ml of ice-cold Buffer A (1 mM HEPES, pH 7.2, 15 mM KCl, 1.5 mM MgAc, 1 mM DTT, and 1X PIC). Cells were then homogenized with 40 strokes of a loose-fitting pestle in a Dounce homogenizer then passed through a 23 G needle 5 times. After homogenization, 500 µl of ice-cold Buffer B (220 mM HEPES, pH 7.2, 375 mM KCl, 22.5 mM MgAc, 1 mM DTT, and 20 µM DNase I) was added and samples were then centrifuged at 750 xg for 10 min. The supernatant was then decanted over a QuadroMACS™ LS column (Miltenyi Biotec, 130-042-976) that had previously been equilibrated with 0.5 % BSA in PBS, and then collected non-lysosomal fraction (Flow) to flow through via gravity. The pellet was subjected to re-addition of 4 ml ice cold Buffer A, 500 µl ice cold Buffer B and then re-suspended and re-centrifuged. This second supernatant was also passed over the column and allowed to flow through via gravity. DNase I (10 µl/ml in PBS) was added and the column was then incubated for 10 min and the washed with 1 ml ice cold PBS. Lysosomes were eluted by removing the column from the magnetic assembly, adding 100 µl of PBS for immunoblotting / M1 buffer (10 mM Tris, pH 7.5, 250 mM Sucrose, 150 mM KCl, 3 mM β-mercaptoethanol, and 20 mM CaCl_2_) for v-ATPase activity and Proton Translocation assay and forced through the column using a plunger. For the co-immunoprecipitation assay, lysosomes were re-suspended in 1X NativePAGE™ sample buffer with 1% Digitonin. After incubation on ice for 30 min, the resolved samples centrifuged at 20,000 xg for 30 min, and then collected lysosomes including solubilized native proteins. Fresh lysosomes (100 µg) were pulled down with C1/6.1 antibody for 24 hr at 4 ºC with gentle rotation.

### v-ATPase activity assay

Lysosome-enriched fractions (fibroblasts: 4 µg; mouse brain: mixture of 20 µl each Optiprep fraction from 15-18) were mixed with 0.052% NaN_3,_ which is an inhibitor of P- and F-type ATPase, to minimize nonspecific ATPase activity. The v-ATPase activity measured using ATPase assay kit according to the manufacturer’s protocol. Control samples were measured in the presence of the v-ATPase inhibitor ConA (1 µM) and the experimental values were subtracted accordingly. Absorbance was measured at 650 nm and solutions of P_i_ were used to generate a standard curve.

### Proton translocation assay

Proton transport activity into the lumen of isolated lysosomes was measured by fluorescence quenching of 9-amino-6-chloro-2-methoxyacridine (ACMA) in the presence or absence of 1 µM ConA. Lysosome-enriched fractions (25 µg) were added to a cuvette containing 2ml of reaction buffer (10 mM BisTrisPropene [BTP]-MES, pH 7.0, 25 mM KCl, 2 mM MgSO_4_, 10 % glycerol and 2 µM ACMA). The reaction was started by the addition of 1 mM ATP in BTP, pH 7.5, a measurement (Ex412/Em480) taken every 5 sec for 600 sec on a SpectraMax M5 multimode reader (Molecular Devices).

### Subcellular fractionation

For fibroblasts, cells were washed and harvested with 1X PBS then re-suspended in ice cold membrane preparation buffer (5 mM Tris, pH 7.4, 250 mM Sucrose, 1 mM EGTA, and 1X PIC). Cells were then briefly homogenized then passed through 26 G needle 10 times. For mouse brain, half of an adult mouse brain was homogenized in 10x volume homogenization buffer (HB: 250 mM sucrose, 10 mM Tris-HCl, pH 7.4, 1 mM EDTA, and protease and phosphatase inhibitors) by 40 strokes in a Teflon-coated pestle. For human brain, the frozen brain tissue (frontal cortex, Brodmann Area 9; supplementary table S2) was homogenized in 10x volume HB by 3 strokes of an automatic homogenizer. After 10 min on ice, samples were then centrifuged at 1,000 xg for 10 min. The supernatants were fractionated into cytosolic and membrane fractions by high-speed centrifugation at 150,000 xg for 60 min. The membrane fractions (pellet after centrifugation) lysis with 100 µl lysis buffer (10 mM Tris, pH 7.4, 150 mM NaCl, 1 mM EDTA, 1 mM EGTA, 1 % Triton X-100, 0.5 % NP-40) and sonication for immunoblotting

### Iodixanol step gradient

Half of an adult mouse brain was homogenized in 10x volume HB by 40 strokes in a Teflon-coated pestle. Lysates were centrifuged at 1,000 xg for 20 min to generate the post nuclear supernatant (PNS). The PNS was then adjusted to 25% OptiPrep with 50% OptiPrep in HB. The resulting mixture, 2 ml in 25% OptiPrep, was placed at the bottom of a clear ultracentrifuge tube and was overlaid successively with 1.5 ml each of 20, 15, 14, 12.5, 10, and 5% OptiPrep in cold HB. The gradients were centrifuged for 18 h at 100,000 xg at 4 °C in a SW 40 rotor (Beckman Coulter). 500 µl fractions were collected from the top of the ultracentrifuge tubes and analyzed by WB analysis.

### Gel electrophoresis and immunoblotting

Samples were mixed with 2X urea sample buffer (9.6 % SDS, 4M Urea, 16% Sucrose, 0.005% Bromophenol blue [BPB], and 4.6% β-mercaptoethanol) and incubated 15 min at 55 ºC for V0 subunits of v-ATPase electrophoresis, otherwise samples mixed with 1X SDS sample buffer (62.5 mM Tris, pH 6.8, 10 % Glycerol, 1 % SDS, 20 mM DTT, 5% β-mercaptoethanol, and 0.005 % BPB) and incubated 5 min at 100 ºC followed by electrophoresis on Novex™ 4-20% Tris-Glycine gradient gels (Invitrogen, WXP42020BOX; WXP42026BOX). Proteins were transferred onto 0.2 µm nitrocellulose membranes (Pall Laboratory, p/n 66485) and the membrane was blocked using 5% non-fat milk or 5% BSA for detecting phospho-proteins. The membrane was then incubated overnight in primary antibody followed by incubation with HRP conjugated secondary antibody. The blot was developed using ECL-kits.

### Co-Immunoprecipitation (co-IP)

Proteins were prepared from cells using ULTRARIPA® kit for Lipid Raft, which can efficiently and rapidly extract membrane proteins/membrane-associated proteins enriched in lipid rafts with native structure and native function, according to the manufacturer’s protocol. 1 mg of cell lysates were pulled down with proper antibodies for 24 hr at 4 ºC with gentle rotation. The mixture was then incubated with 30 µl of Protein A/G Mix Magnetic Beads and washed with lysis buffer and Magnetic Stand. The samples were dissociated by 2X Urea sample buffer. For the peptide competition assay, 500 µg of fresh cell lysates was pulled down with C1/6.1 antibody in presence of varying amounts of each peptide for 24 hr at 4 ºC with gentle rotation.

### Native gel immunoblot analysis

The pellet from the cell membrane fraction was resolved with 1% Digitonin and 1X NativePAGE™ sample buffer, incubated on ice for 30 min, and then centrifuged at 20,000 xg for 30 min. 15 µg of sample was mixed with 1 % NativePAGE™ G-250 sample additive followed by electrophoresis on NativePAGE™ Novex^®^ 4-16% Bis-Tris Gels in 1X NativePAGE™ Running Buffer then transfer using 1X NuPAGE® Transfer buffer and 0.45 µm PVDF membranes (Millipore, IPVH00010) according to manufacturer’s protocol.

### Confocal fluorescence microscopy

Cells were fixed with 4 % PFA for 20 min, blocked with 5 % horse serum, incubated with anti-cathepsin B goat pAb (CTSB; 1/250) for lysosomal marker and anti-V1D rabbit mAb (1/100) overnight at 4 ºC, followed by fluorescence tagged secondary antibodies; Donkey anti-Goat Alexa Fluor 488 (1/500) and goat anti-Rabbit Alexa Fluor 568 (1/500). Cells were imaged using a plan-Apochromat 40x/1.4 oil DIC objective lens on the laser scanning confocal microscope, LSM 510 META, with LSM software v3.5 (Carl Zeiss MicroImage Inc). Images were analyzed using ImageJ program (NIH). Co-localization was determined using the Coloc 2 analysis tool on ImageJ software.

### Proximity ligation assay

Proximity ligation assays were performed according to the manufacturer’s protocol using the following reagents in the Duolink^®^ In Situ Detection Reagents Red kit. In brief, cells were plated on cover slips at a concentration of 10^5^/ml. Cells were fixed using 4 % PFA in PBS for 15 min, then permeabilized using 0.01 % Triton in PBS for 20 min at room temperature. All subsequent incubations were performed at 37 ºC in a humid chamber. Non-specific binding was minimized with the manufacturer’s blocking buffer for 30 min. Cells were incubated with primary anti-APP (C1/6.1) mouse (1/100) and anti-V0a1 rabbit mAb (1/100) overnight at 4 ºC and washed with buffer A (twice, 5 min). Coverslips were incubated with ligation mix for 30 min, washed twice with buffer A, and incubated with amplification mix. Coverslips were then washed with provided Buffer B twice (10 min), followed by 1 min washing with 0.01x Wash Buffer B. The samples were mounted with Duolink® In Situ Mounting Medium with DAPI in the dark for 15 min at room temperature. The images were taken using a Zeiss LSM880 laser scanning confocal microscope at 20x objective lens. The number of signals per cell were analyzed using ImageJ program and then normalized to the protein levels.

### Mass spectrometry (MS) analysis

The stained protein gel regions (fig. S2A, Bands 1-6) were excised, and in-gel digestion was performed overnight with mass spectrometry grade Trypsin at 5 ng/µl in 50 mM NH_4_HCO_3_ digest buffer. After acidification with 10 % formic acid, peptides were extracted with 5 % formic acid/50 % acetonitrile (v/v) and concentrated to a small droplet using vacuum centrifugation. Desalting of peptides was done using hand packed SPE Empore C18 Extraction Disks (aka Stage Tips, 3M. St. Paul, MN, USA) as described (*95*). Desalted peptides were again concentrated and reconstituted in 10 µl 0.1 % formic acid in water. An aliquot of the peptides was analyzed by nano-LC-MS/MS using an Easy nLC 1,000 equipped with a self-packed 75 µm × 20 cm reverse phase column (ReproSil-Pur C18, 3 m, Dr. Maisch GmbH, Germany) coupled online to QExactive HF Orbitrap mass spectrometer via a Nanospray Flex source (all instruments from Thermo Scienctific, Waltham, MA, USA). Analytical column temperature was maintained at 50 ºC by a column oven (Sonation GmBH, Germany). Peptides were eluted with a 3-40% acetonitrile gradient over 60 min at a flow rate of 250 nl/min. The mass spectrometer was operated in the DDA mode with survey scans acquired at a resolution of 120,000 (at m/z 200) over a scan range of 300-1,750 m/z. Up to 15 of the most fragmentation by higher-energy collisional dissociation with a normalized collision energy of 27. The maximum injection times for the survey and MS/MS scans were 60 ms. And the ion target value for both scan modes was set to 3e6. All acquired mass spectra were first converted to mascot generic format files (mgf) using Proteome Discoverer 1.4 and generated mgf files searched against a human protein database (SwissProt, 20, 210 sequences, 2014) using Mascot (Matrix Science, London, UK; version 2.7.0 www.matrixscience.com). Decoy proteins are added to the search to allow for the calculation of false discovery rates (FDR). The search parameters are as follows: (i) two missed cleavage tryptic sites are allowed; (ii) precursor ion mass tolerance = 10 ppm; (iii) fragment ion mass tolerance = 0.3 Da; and (iv) variable protein modifications are allowed for methionine oxidation, deamidation of asparagine and glutamines, cysteine acrylamide derivatization and protein N-terminal acetylation. MudPit scoring is typically applied using significance threshold score *p*<0.01. Decoy database search is always activated and, in general, for merged LS-MS/MS analysis of a gel lane with *p*<0.01, false discovery rate for protein ID averaged around 1 %.

### Protein preparation for Glide molecular docking

The cryo-EM structure of the v-ATPase was retrieved from the Protein Data Bank (PDB ID: 6VQ7), and it was refined with Protein Preparation Wizard implemented in Maestro 12. The protein structure was imported into workspace and preprocessed to assign bond orders, add hydrogen atoms, create zero-order bonds to metals, create disulfide bonds, and to delete water molecules beyond 5 Å from hetero groups. In addition, missing atoms in residues and missing loops were added using Prime to generate a complete protein structure (note: the 34 missing amino acids (R133 – G169) in the V0a1 subunit was not built using Prime due to the size of the missing loop). The protein structure was further refined via automated H-bond assignment and restrained minimization with OPLS 2005 force field by converging heavy atoms to 0.3 Å RMSD. The N-terminal domain of the V0a1 subunit (E2 – Y363) was retrieved by deleting other subunits. The missing loop (R133 – G169) was built using the 3D builder module in Maestro 12, and the complete N-terminal domain of the V0a1 subunit was further refined with Protein Preparation Wizard. The folding states of the missing loop were searched by MD simulation using the Amber Molecular Dynamics Package. The truncated protein structure (N111 – V271) with ACE and NME terminal caps was solvated with the Amber ff14SB force field and TIP3P explicit water model in an octahedron periodic box using a buffer distance of 14.0 Å containing 150 mM NaCl in 20,827 water molecules. The solvated protein structure was treated to 3 consecutive minimization stages: (i) 1,000 steps of steepest descent and 1,000 steps of conjugate gradient minimization with 100 kcal mol^-1^ Å^-2^ restraints on all atoms except water molecules; (ii) 2,500 steps of steepest descent and 2,500 steps of conjugate gradient minimization without restraints; and (iii) 1,000 steps of steepest descent and 2,500 steps of conjugate minimization with 100 kcal mol^-1^ Å^-2^ restraints on N111 – F130 and Q252 – V271, which are helix units connected to the rest of the V0a1 subunit. After minimization, equilibration of the solvated protein was performed at 303.15 K for 100 ns under constant volume, with a timestep of 2 fs, and with SHAKE algorithm employed to restrain on calculation of forces of bonds containing hydrogen atoms. In the equilibration, amide backbones of N112 – K129 and F170 – V271 were fixed with 10 kcal mol^-1^ Å^-2^ restraints. The last frame of the MD equilibration was extracted from the simulation trajectories using the CPPTRAJ program in the AmberTools20, and it was subjected to the Glide docking after removing water molecules and ions.

### Glide molecular docking and MD simulations

The last frame of the MD simulation was refined by Protein Preparation Wizard, and a protein grid for Glide dock was generated. Van der Waals radius was scaled by decreasing the default value of scaling factor to 0.8 to soften the potential for nonpolar parts of the receptor. The length of the inner box was increased to 20 Å^3^ from the default value (10 Å^3^) to explore all available binding sites of the target protein in the molecular docking. A ligand structure (NGpYEN) was built using the 3D build module and the N- and C-terminal amino acids were capped with the acetyl group (ACE) and N-methylamine (NMA) in the refinement step, respectively. Glide software (v 8.7) in Maestro 12 was utilized to dock the ligand structure to the protein grid in the standard precision mode with the OPLS 2005 force field. The ligand structure was flexibly docked by sampling rotatable bonds, nitrogen inversions, and ring conformations. For the output of Glide docking, 200 poses were included for post-docking minimization, and 95 binding poses of the NGpYEN structure were obtained. After visual analysis by considering docking scores, 16 diverse binding poses of the NGpYEN structure at multiple binding sites were selected for further modeling with molecular dynamics. Each complex structure of the truncated V0a1 with each binding pose of NGpYEN was solvated with TIP3P explicit water model in an octahedron periodic box using a buffer distance of 14.0 Å containing 150 mM NaCl in ca. 13,000 water molecules. Amber ff14SB and phosaa14SB force fields were applied to the solvated protein – ligand complex for MD simulation. Minimization and equilibration of the solvated protein – ligand complexes were conducted for 10 ns with the same parameters and constraints aforementioned. The last frame of the equilibration was obtained from the trajectories, and additional amino acids were added to the ligand structure in Maestro 12. Multiple minimization and equilibration were iteratively conducted for 480 ns with the same parameters and various constraints. VMD software was used to visually analyze the trajectories, and the binding pose of the ligand that showed similar binding pattern to the known structure of pY to its pocket (fig. S7A) was selected. Figures were generated using Pymol (The PyMOL Molecular Graphics System, Version 2.0 Schrodinger, LLC).

### Estimation of Binding Affinity using MM-GBSA

The binding free energies of the V0a1 model structure in complex with phosphorylated and unphosphorylated ligands were estimated by generalized Born surface area continuum solvation method (MM-GBSA). The truncated V0a1 – a phosphorylated ligand (ACE-MQQNGpYENPTYK-NMA) structure was obtained from the previous restricted MD simulation followed by protein preparation in Maestro 12. The model structure without the phosphorylated Y was obtained by mutating pY to Y in Maestro 12. These model structures were solvated as described above, and the whole systems were minimized by 2,500 steps of steepest descent and 2,500 steps of conjugate gradient minimization without restraints. After minimization, the systems were heated from 0 K to 303.15 K for 300 ps and equilibrated for 100 ps under NVT condition followed by another equilibration for 100 ps under NPT condition. Finally, 10 ns production run for each system was executed at a constant pressure (1 bar) and a constant temperature (303.15 K). The estimated binding free energy of each system was calculated with all trajectories (1,000 frames) of the complexes.

### Quantification and statistical analysis

All quantitative data were subjected to two-tailed unpaired Student’s *t*-test for single comparison, or one-way ANOVA analysis for multiple comparisons with Sidak’s analysis using GraphPad Prism 9. For cellular data, 3 or more biological replicates with mean ± SEM were represented in bar graph with individual dots. Differences were considered significant with P <0.05. Statistical parameters, including the definitions and value of sample size (n), deviations and P values, are reported in the corresponding figure legends. Statistically significance is represented with asterisks or hatch marks * or ^#^, *p* ≤ 0.05; ** or ^##^, *p* ≤ 0.005; *** or ^###^, *p* ≤ 0.0005; **** or ^####^, *p* ≤ 0.0001.

## Acknowledgments

We thank Dr. Kuglae Kim (Yonsei University, Republic of Korea) for his advice on structural analyses. We are very grateful to Dr. Efrat Levy (Nathan S. Kline Institute; NYU Grossman School of Medicine) for supervising animal breeding, and thank Dr. Monica Pawlik, Chunfeng Huo, and Steven DeRosa (Nathan S. Kline Institute) for maintaining animals. Human tissues were obtained from the Brain Bank for Developmental Disabilities and Aging.

## Author contributions

Conceptualization: EI, RAN, YJ

Methodology: EI, RAN, JYC, HEB, TAN, JW

Investigation: EI, PS, SD, HEB, TAN, JYC

Visualization: EI

Supervision: RAN

Writing—original draft: EI, RAN,

Writing—review & editing: EI, RAN, YJ, JHL, PS, JYC

## Data and materials availability

Materials generated in this study are available and may require completion of a material transfer agreement. All data are available in the main text or the supplementary materials.

