## Supplementary Materials for "Lysosomal dysfunction in Down Syndrome and Alzheimer mouse models is caused by selective v-ATPase inhibition by Tyr^682^ phosphorylated APP βCTF"

Eunju Im et al.

**This PDF file includes:**

Figs. S1 to S10
Tables S1 to S2

Supplementary Figure S1_related to Figure 1

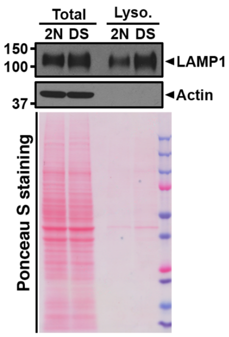


**Fig. S1. Western blot analysis to confirm purity of lysosome isolation (related to Fig. 1)**.

(**A**) Cell lysates (Total) or lysosome enriched fraction (Lyso) of 2 yr 2N and DS fibroblasts were immunoblotting with anti LAMP1 and Actin. LAMP1 served as a lysosomal marker and Actin served as a loading control. The Ponceau S staining represent the amounts of loaded proteins.

**Supplementary Figure S2_related to Figure 1**

A

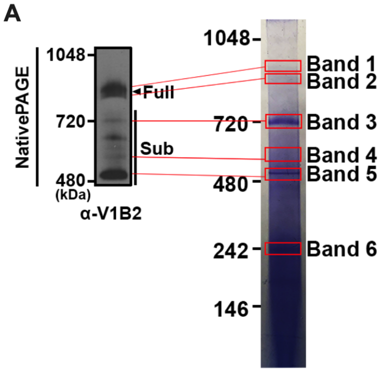


B

| Alternate ID | Mr/Da | Full complex | | Sub complex | | | Negative |
|---|---|---|---|---|---|---|---|
| | | Band 1 | Band 2 | Band 3 | Band 4 | Band 5 | Band 6 |
| | | Total Spectral Counts | | | | | |
| ATP6V0a1 | 96 kDa | 7 | 11 | 1 | 20 | | |
| ATP6V0a2 | 98 kDa | | 5 | 4 | 13 | | |
| ATP6V0d1 | 40 kDa | 7 | 8 | 4 | 9 | | |
| ATP6V1A | 68 kDa | 23 | 34 | 10 | 13 | 13 | |
| ATP6V1B2 | 57 kDa | 22 | 29 | 7 | 11 | 6 | |
| ATP6V1C1 | 44 kDa | 13 | 15 | 1 | | | |
| ATP6V1D | 28 kDa | 6 | 11 | | 2 | 2 | |
| ATP6V1E1 | 26 kDa | 12 | 21 | 1 | 2 | | |
| ATP6V1F | 13 kDa | 1 | 7 | 1 | | | |
| ATP6V1G1 | 14 kDa | 5 | 7 | 1 | | | |
| ATP6V1H | 56 kDa | 6 | | | | | |
| ATP6AP1 | 52 kDa | 1 | 1 | 1 | 8 | 2 | |
| ATP6AP2 | 39 kDa | | | 6 | 9 | 5 | |

**Fig. S2. Mass spectrometry analysis to identify the bands matching with the detected bands with a V1B2 subunit antibody (related to fig. 1).**

(**A**) *Left panel:* Membrane fractions were resolved using the native PAGE and immunoblotted with anti-V1B2 antibody. *Right panel:* Membrane fractions were resolved using native PAGE (same blot with Fig. 1E [DS]) and fixed with fix solution (40% methanol, 10% acetic acid). The fixed gel stained with Coomassie R-250 Stain (0.02% Coomassie R-250 in 30% methanol and 10% acetic acid) and then destained with destaining solution (8% acetic acid). (**B**) The table represents total mass spectrometry spectral counts of each band to indicate relative abundance of the v-ATPase subunits.

### Supplementary Figure S3_related to Figure 1

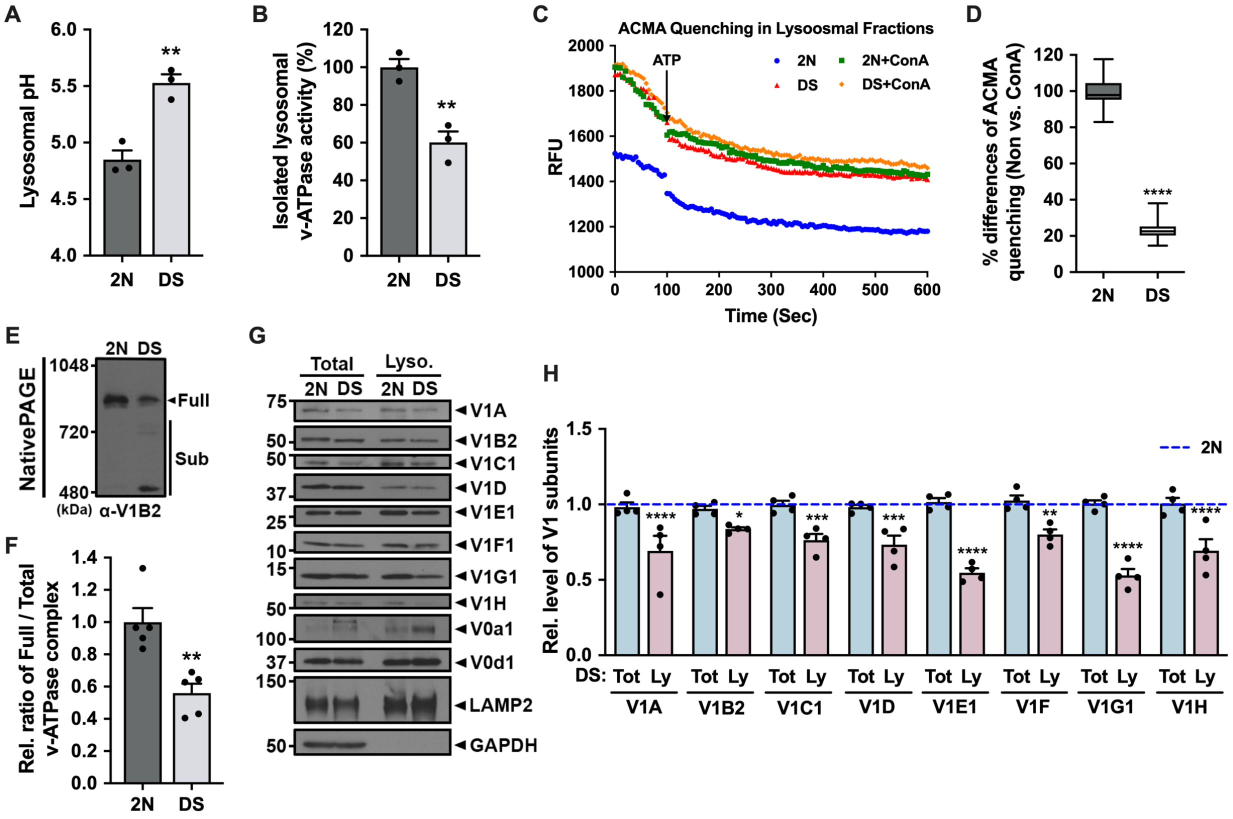


**Fig. S3. Assembly of the lysosomal v-ATPase complex is impaired in 5-month old DS fibroblasts (related to Fig. 1).**

(**A**) Lysosomal pH values measured ratiometrically using LysoSensor Yellow/Blue (Y/B) dextran using lysosome enrichment fraction from 5 months fibroblasts (n=3, three independent, duplicate). (**B**) Lysosomal v-ATPase activity measured colorimetrically as ATP hydrolysis with and without a v-ATPase inhibitor (Concanamycin A; ConA). Activity assay is performed on lysosomal fractions pre-treated with inhibitors of P- and F-type ATPases (*o*-vanadate) to minimize nonspecific ATPase activity using 5 months fibroblasts (n=3, three independent, triplicate). (**C, D**) Lysosomal v-ATPase activity measured fluorometrically using a pH gradient probe (ACMA method) to measure $H^+$ transport into lysosomes by the v-ATPase with and without a ConA using 5 months fibroblasts (n=3, three independent). (**E**) Membrane fractions from 5 months fibroblasts were resolved using native PAGE and immunoblotted with anti-V1B2 antibody. (**F**) The graph represents relative ratio of full complex divided by total (full plus sub complexes) complexes of v-ATPase (n=5, five independent). (**G**) Immunoblot of v-ATPase subunit distributions in total lysates (Total) and lysosomal enriched (Lyso.) fraction of 5 months 2N and DS fibroblasts. LAMP2 served as a marker for lysosome and GAPDH served as a loading control for total lysates. (**H**) The graphs show band intensity of each v-ATPase subunits (n=4, four independent). Quantitative data (A, B, D, F, and H) are presented as mean values with ±SEM, two-tailed unpaired t test (A, B, D, and F), ordinary two-way ANOVA with Šidák's multiple comparisons test (H); *, $P < 0.05$; **, $P < 0.005$; ***, $P < 0.0005$; ****, $P < 0.0001$. Each dot represents average value of technical replicates from each independent experiment.

Supplementary Figure S4_related to Figure 3

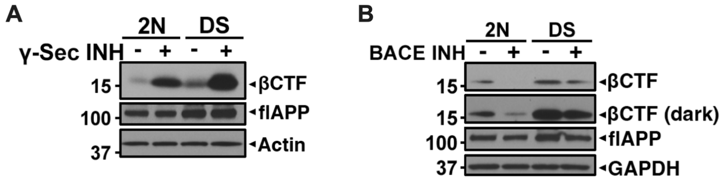


**Fig. S4. Western blot analysis to confirm the effect of secretase inhibitors (related to Fig. 3).**

(**A, B**) The immunoblot represents βCTF and flAPP protein levels in total lysates from 2yr 2N and DS fibroblast after treated with either DMSO (-), γ-Sec INH (10 μM) (*A*), or BACE INH (10 μM) (*B*) for 24 hr. Cell lysates were immunoblotted with anti-APP (6E10) antibody. Actin and GAPDH served as a loading control.

Supplementary Figure S5_related to Figure 3

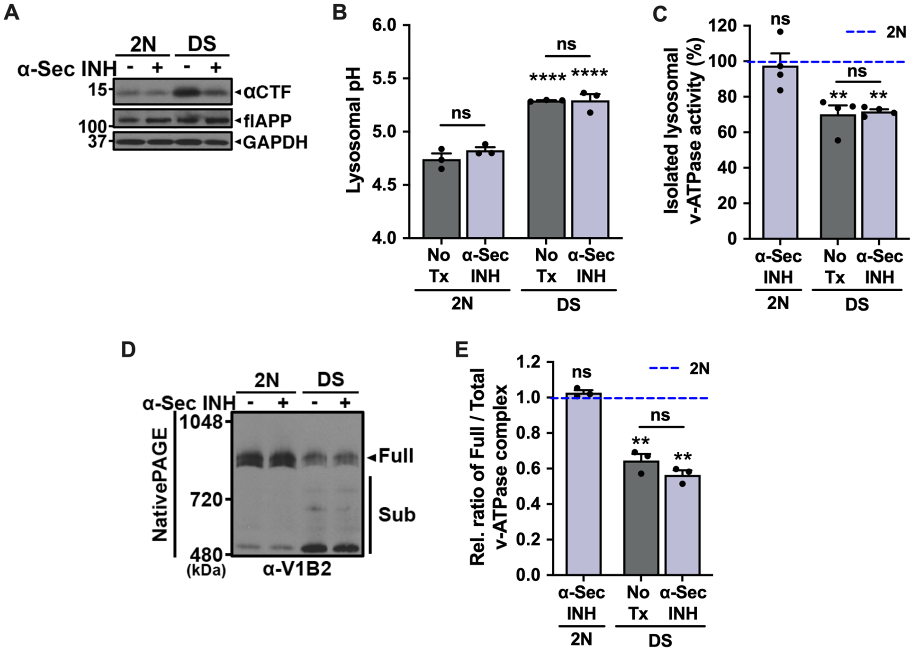


**Fig. S5. Lysosomal v-ATPase activity is not affected by APP-αCTF (related to Fig. 3).**

(**A**) The immunoblot represents αCTF and flAPP protein levels in total lysates from 2yr 2N and DS fibroblast after treated with either DMSO (-) or α-Sec INH (20 µM) for 24 hr. Cell lysates were immunoblotted with anti-APP (C1/6.1) antibody. GAPDH served as a loading control. (**B**) Lysosomal pH of 2N and DS fibroblasts treated with either DMSO (No Tx) or α-secretase inhibitor, TAPI-1 (α-Sec INH; 20 µM), for 24 hr determined by LysoSensor Y/B dextran (n=3, three independent, triplicate). (**C**) Lysosomal v-ATPase activity measured colorimetrically as ATP hydrolysis with and without a ConA. Activity assay performed on lysosomal fractions from 2N and DS fibroblast after treated with DMSO or 20 µM α-Sec INH for 24 hr, pre-treated with *o*-vanadate to minimize nonspecific ATPase activity (n=4, four independent, duplicate). (**D**) Membrane fractions from 2N and DS fibroblasts treated with DMSO (-) or α-Sec INH (+) were resolved using the native PAGE and immunoblotted with anti-V1B2 antibody. (**E**) The graph represents relative ratio of full complex divided by total (full plus sub complexes) complexes of v-ATPase (n=3, three independent). Quantitative data (B, C, and E) are presented as mean values with ±SEM, ordinary two-way ANOVA with Šidák's multiple comparisons test; **, $P < 0.005$; ****, $P < 0.0001$; ns, not significant. Each dot represents average value of technical replicates from each independent experiment.

Supplementary Figure S6_related to Figure 4

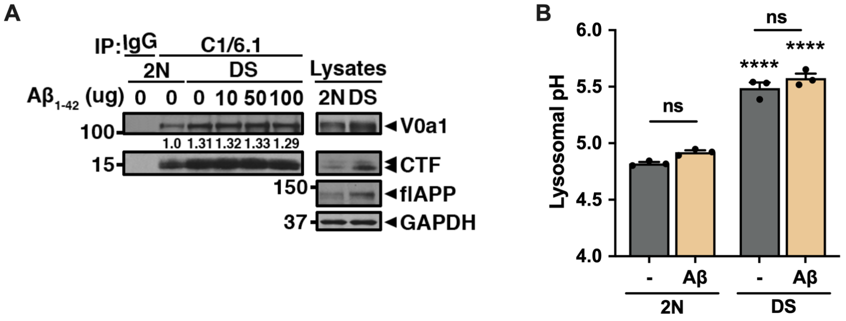


**Fig. S6. A$\beta_{1\text{-}42}$ peptide is no effect on v-ATPase and lysosomal pH (related to Fig. 4).**

(**A**) Cell lysates of 2yr 2N and DS fibroblast were incubated with A$\beta_{1\text{-}42}$ peptide following indicated amount for 24 hr, then immunoprecipitated with anti-APP (C1/6.1) antibody, followed by immunoblotting with anti-V0a1 antibody. The values at the bottom of the left IP blot indicate the relative intensities of IP-ed V0a1normalized by IP-ed APP-CTFs. The quantification in this experiment is confirmed in two other experiments. (**B**) Lysosomal pH of 2yr 2N and DS fibroblasts treated with either DMSO (-) or 5 μM A$\beta_{1\text{-}42}$ peptide for 24 hr determined by LysoSensor Y/B dextran (n=3, three independent, triplicate). Quantitative data (B) is presented as mean values with ±SEM, ordinary two-way ANOVA with Šidák's multiple comparisons test; ****, $P < 0.0001$; ns, not significant. Each dot represents average value of technical replicates from each independent experiment.

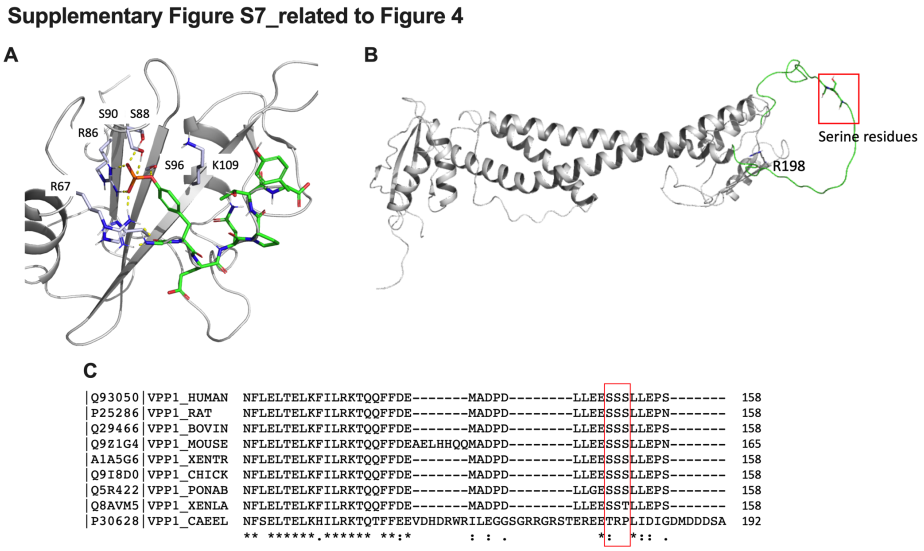


**Fig. S7. Structure analysis to find the binding region in the V0a1 subunit (related to Fig. 4).**

(**A**) A known binding site of phosphorylated tyrosine residue (X-ray co-crystal structure of Grb2-SH2 (gray) and AICD (green) complex, PDB ID: 3MXC). The phosphate unit shows hydrogen bond and salt bridge interactions as presented with yellow dashed lines. The phenyl ring of tyrosine form $\pi$ – hydrophobic contacts with the methylene unit of K109. (**B**) The cryo-EM structure of V0a1 subunit of v-ATPase (PDB ID: 6VQ7) and the unsolved loop. An unsolved loop in the cryo-EM structure is built by using maestro molecular modeling software (green), and three consecutive serine residues (red box) in the unsolved loop are labeled. (**C**) Multiple sequence alignment of the unsolved loop of V0a1 subunits shows that three serine residues (red box) are highly conserved among species. The first column shows UniProt entry code, and VPP1_HUMAN, RAT, BOVIN, MOUSE, XENTR, CHICK, PONAB, XENLA, and CAEEL represent human, rat, bovine, mouse, frog, chicken, orangutan, C. elegans v-ATPase V0a1 subunits, respectively.

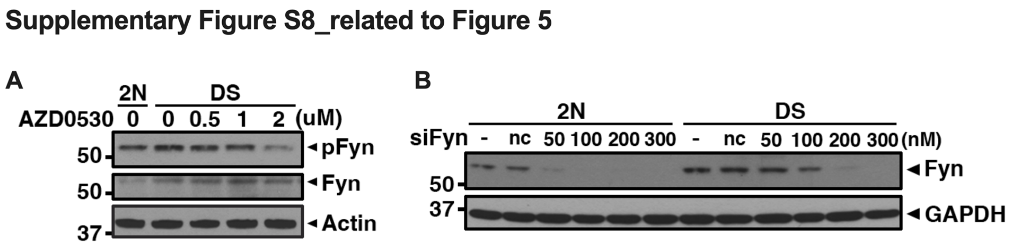


**Fig. S8. Western blot analysis to adjust concentrations of inhibitor or siRNA (related to Fig. 5)**

(**A**) Immunoblot represents phospho-Fyn (pFyn) and Fyn kinase protein levels in 2yr 2N and DS fibroblast after treated with DMSO (-) or AZD0530 following indicated amount for 24 hr. (**B**) Immunoblot represents Fyn kinase protein levels in 2yr 2N and DS fibroblast after transfected without (-) or with siNC or siFyn following indicated amount for 48 hr.

Supplementary Figure S9_related to Figure 6

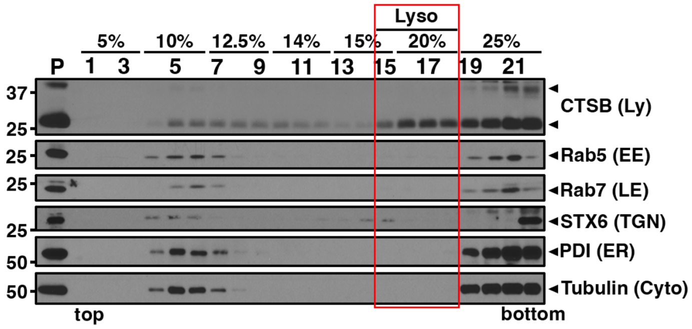


**Fig. S9. Western blot analysis of fractions from the iodixanol step gradient (related to Fig. 6)**

Adult mouse brain homogenates were fractionated through an iodixanol step gradient. Immunoblot analysis showed the distribution of cathepsin B (CTSB; lysosome marker; Ly), Rab5 (early endosome marker; EE), Rab7 (late endosome marker; LE), syntaxin 6 (STX6; trans-golgi network marker; TGN), PDI (endoplasmic reticulum marker; ER), and Tubulin (Cytosolic marker; Cyto).

**Supplementary Figure S10_related to Figure 6**

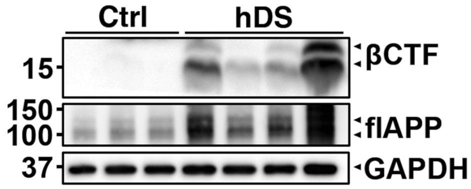


**Fig. S10. Protein levels of βCTFs and flAPP increased in the homogenate from the frozen brain of human DS (related to Fig. 6).**

Immunoblot represents CTFs and flAPP protein levels in frozen frontal cortex (BA 9) from control and DS subjects. Brain homogenates were immunoblotted with anti-APP (6E10) antibody. GAPDH served as a loading control.

**Table S1. Key resource**

| REAGENT or RESOURCE | SOURCE | IDENTIFIER |
| --- | --- | --- |
| Antibodies | | |
| Mouse anti-APP (C1/6.1) | Mathew et al., 2002 | |
| Mouse anti-APP (6E10) | BioLegend | Cat# 803001; RRID: AB_2564653 |
| Rabbit anti-APP (Phospho-Tyr$^{682}$) | GenScript | Cat# A00697-40; RRID: AB_1108125 |
| Rabbit anti-V0a1 | Abcam | Cat# ab176858; RRID: AB_2802122 |
| Rabbit anti-V0b | Abcam | Cat# ab107189; RRID: AB_10865216 |
| Rabbit anti-V0c | Abcam | Cat# ab104374; RRID: AB_10712593 |
| Mouse anti-V0d1 | Abcam | Cat# ab56441; RRID: AB_940402 |
| Rabbit anti-V0e2 | Abcam | Cat# ab178934; RRID: AB_2892774 |
| Rabbit anti-V1A | GeneTex | Cat# GTX110815; RRID: AB_1949704 |
| Rabbit anti-V1B2 | Abcam | ab73404; RRID: AB_1924799 |
| Rabbit anti-V1C1 | Abcam | Cat# ab87163; RRID: AB_1951504 |
| Rabbit anti-V1D | Abcam | Cat# ab157458; RRID: AB_2732041 |
| Rabbit anti-V1E1 | Abcam | Cat# ab111733; RRID: AB_10861729 |
| Rabbit anti-V1F | Abcam | ab190789; RRID: AB_2892713 |

| | | |
|---|---|---|
| Rabbit anti-V1G1 | Abcam | Cat# ab174243; RRID: AB_2892716 |
| Rabbit anti-V1H | Abcam | Cat# ab96120; RRID: AB_10679342 |
| Rabbit anti-Fyn | Cell Signaling Technology | Cat# 4023; RRID: AB_10698604 |
| Rabbit anti-phospho-Fyn (Tyr420) | MyBioSouce | Cat# MBS9128730; RRID: AB_2892719 |
| Mouse anti-LMAP1 | Developmental Studies Hybridoma Bank | Cat# H4A3; RRID: AB_2296838 |
| Mouse anti-LMAP2 | Developmental Studies Hybridoma Bank | Cat# H4B4; RRID: AB_2134755 |
| Goat anti-cathepsin B | Neuromics | Cat# GT15047; RRID: AB_2737184 |
| Rabbit anti-cathepsin D | Lee et al., 2015; 2022 | |
| Rabbit anti-Rab5 | Abcam | Cat# ab218624; RRID: AB_2892717 |
| Mouse anti-Rab7 | Abcam | Cat# ab50533; RRID: AB_882241 |
| Mouse anti-PDI | GeneTex | Cat# GTX22792; RRID: AB_384853 |
| Rabbit anti-Syntaxin 6 | Cell Signaling Technology | Cat# 2869; RRID: AB_2196500 |
| Mouse anti-Tubulin | Sigma-Aldrich | Cat# T8535; RRID: AB_261795 |
| Mouse anti-Actin | Sigma-Aldrich | Cat# A1978; RRID: AB_476692 |
| Mouse anti-GAPDH | Millipore Sigma | Cat# CB1001; RRID: AB_2107426 |
| Rabbit anti-Na, K-ATPase alpha | Cell Signaling Technology | Cat# 3010; RRID: AB_2060983 |

| | | |
|---|---|---|
| Mouse Normal IgG control | Millipore | Cat# 12-371; RRID: AB_145840 |
| Rabbit Normal IgG control | Millipore | Cat# 12-370; RRID: AB_145841 |
| Peroxidase AffiniPure Donkey Anti-Mouse IgG (H+L) | Jackson ImmunoResearch | Cat# 715-035-150; RRID: AB_2340770 |
| Peroxidase AffiniPure Donkey Anti-Rabbit IgG (H+L) | Jackson ImmunoResearch | Cat# 711-035-152; RRID: AB_10015282 |
| Peroxidase AffiniPure Donkey Anti-Goat IgG (H+L) | Jackson ImmunoResearch | Cat# 705-035-147; RRID: AB_2313587 |
| VeriBlot for IP detection Reagent (HRP) | Abcam | Cat# ab131366; RRID: AB_2892718 |
| Donkey anti-Goat Alexa Fluor 488 | Molecular Probes | Cat# A-11055; RRID: AB_2534102 |
| Goat anti-Rabbit Alexa Fluor 568 | Molecular Probes | Cat# A-11036; RRID: AB_10563566 |
| Chemicals, peptides, and recombinant proteins | | |
| L-685, 459 | Tocris Bioscience | Cat# 2627; CAS: 292632-98-5 |
| β-Secretase inhibitor IV | Calbiochem | Cat# 565788; CAS: 797035-11-1 |
| TAPI-1 | Calbiochem | Cat# 579051; CAS: 171235-71-5 |
| Saracatinib (AZD0530) | Selleck Chemical LLC | Cat# S1006; CAS: 379231-04-6 |
| Concanamycin A (ConA) | Sigma-Aldrich | Cat# C9795; CAS: 80890-47-7 |
| DNase I | Sigma-Aldrich | Cat# 10104159001 |
| Halt$^{TM}$ Protease and Phosphatase inhibitor Cocktail (100X) | Thermo Fisher Scientific | Cat# 78444 |

| | | |
|---|---|---|
| LysoSensor™ Yellow/Blue dextran | Molecular Probes | Cat# L22460 |
| Dextran conjugated magnetite | Liquid Research LLC | Cat# DexoMAG™40 |
| 9-amino-6-chloro-2-methoxyacridine (ACMA) | Sigma-Aldrich | Cat# A5806; CAS: 3548-09-2 |
| OptiPrep™ Density Gradient Medium | Sigma-Aldrich | Cat# D1556; CAS: 92339-11-2 |
| ECL Chemiluminescent Substrate Reagent | Invitrogen | Cat# WP20005 |
| Immobilon Western Chemiluminescent HRP Substrate | Millipore | Cat# WBKLS0500 |
| PureProteome™ Protein A/G Mix Magnetic Beads | Millipore | Cat# LSKMAGAG10 |
| Precision Plus Protein Dual Color Standards | BIO-RAD | Cat# 161-0374 |
| NativeMark™ Unstained Protein Standard | Thermo Fisher Scientific | Cat# LC0725 |
| Digitonin | Sigma-Aldrich | Cat# D141; CAS: 11024-24-1 |
| NativePAGE™ sample buffer (4X) | Invitrogen | Cat# BN2003 |
| NativePAGE™ G-250 sample additive | Invitrogen | Cat# BN2004 |
| NativePAGE™ Running Buffer (20X) | Invitrogen | Cat# BN2001 |
| NuPAGE™ Transfer Buffer (20X) | Invitrogen | Cat# NP00061 |
| 32% Paraformaldehyde (PFA) | Electron Microscopy Sciences | Cat# 50-980-494; CAS: 30525-89-4 |
| Duolink® In Situ Red Starter Kit Mouse/Rabbit | Sigma-Aldrich | Cat# DUO92101-1KIT |
| Lipofectamine™ RNAiMAX Transfection | Invitrogen | Cat# 13778150 |
| Poly-D-lysine | Sigma | Cat# P0899 |

| | | |
|---|---|---|
| MEM | Gibco | Cat# 10370088 |
| Opti-MEM | Gibco | Cat# 51985034 |
| Fetal Bovine Serum | Gibco | Cat# 16000069 |
| Horse Serum | Gibco | Cat# 16050-122 |
| Penicillin-Streptomycin (10,000 U/ml) | Gibco | Cat# 15140122 |
| Trypsin-EDTA (0.05%), phenol red | Gibco | Cat# 25300120 |
| QNGYENPTY | GenScript | Custom-designed |
| QNG(p)YENPTY | GenScript | Custom-designed |
| QNG(p)YENVTY | GenScript | Custom-designed |
| Amyloid $\beta_{1\text{-}42}$ peptide | GenScript | Cat# RP10034-1 |
| Trypsin gold | Progema | Cat# V5280 |
| SPE Empore C18 Extraction Disks | Aka Stage Tips | Cat# 2215 |
| Self-packed 75µm x 2 cm reverse phase column; Packing: ReproSil-Pur C18-AQ 3µm | Ammerbuch-Entringen | Cat# r13.aq |
| Empty PicoTip Emitter | Now Objective Inc | Cat# PF360-75-10-N5 |
| QExactive HF Orbitrap mass spectrometer | Thermo Fisher Scientific | Cat# 05000L-05999L |
| Nonospray Flex Source | Thermo Fisher Scientific | Cat# ES071 |
| Easy-nLC 1000 nanoflow liquid chromatography (LC) system | Thermo Fisher Scientific | Cat# LC120 |
| Critical commercial assays | | |
| ATPase activity Kit (Colorimetric) | Innova Biosciences (Current: Abcam) | Cat# 601-0120 (Cat# ab270551) |
| ULTRARIPA® kit for Lipid Raft | Biodynamics Laboratory Inc | Cat# F015 |
| Deposited data | | |
| Mammalian v-ATPase structure from rat brain | | PDB: 6VQ7 |

| | | |
|---|---|---|
| Structure of Grb2-SH2 domain and AICD peptide complexes | | PDB: 3MXC |
| Experimental models: Cell lines | | |
| Fibroblast from skin, Thorax/abdomen, Trisomy21, 2 YR | Coriell Cell Repositories | AG06922 |
| Fibroblast from skin, Foreskin, apparently healthy individual, 2 YR | | AG07095 |
| Fibroblast from skin, Thorax, Trisomy21, 5 MO | Coriell Cell Repositories | AG07096 |
| Fibroblast from skin, Foreskin, apparently healthy individual, 5 MO | Coriell Cell Repositories | GM08680 |
| Experimental models: Organisms/strains | | |
| Mouse: Ts2 | Jiang et al., 2019 | |
| Mouse: TRGL6 | Lee et al., 2019 | |
| Mouse: TRGL6 x Ts2 | This study | |
| Mouse: Tg2576 | Lee et al., 2019 | |
| Mouse: 5xFAD | Lee et al., 2019 | |
| Oligonucleotides | | |
| siRNA against human APP: ACUAGUGCAUGAAUAGAUUCUCU CC UUUGAUCACGUACUUAUCUAAGA GAGG | Integrated DNA Technologies | |
| siRNA against human Fyn: GGACUCACCGUCUUUGGAGtt ttCCUGAGUGGCAGAAACCUC | Life Technologies Corp. | Silencer Pre-Designed siRNA ID 1442, Cat# AM51331 |

| | | |
|---|---|---|
| Negative control DsiRNA:CGUUAAUCGCGUAUAAUACGCGUAT AUACGCGUAUUAUACGCGAUUAACGAC | Integrated DNA Technologies | Cat# 51-01-14-03 |
| Software and algorithms | | |
| FIJI | NIH | https://hpc.nih.gov/apps/Fiji.html |
| Photoshop 2021 | Adobe | |
| GraphPad Prism 9 | GraphPad | https://www.graphpad.com/scientific-software/prism/ |
| Maestro 12 | Schrödinger | https://www.schrodinger.com/products/maestro |
| The PyMOL Molecular Graphics System v2.0 | Schrödinger | https://www.schrodinger.com/products/pymol |

**Table S2. Demographic data on the brains in this study**

| No | Group | Age (y) | Gender | PMI (h) |
|---|---|---|---|---|
| 1 | DS | 51 | M | 2 |
| 2 | DS | 55 | M | 5.35 |
| 3 | DS | 56 | M | 4 |
| 4 | DS | 59 | M | 3.4 |
| 5 | DS | 60 | F | 4 |
| 6 | DS | 59 | F | 6 |
| 7 | DS | 63 | M | 14 |
| 8 | DS | 67 | M | 8 |
| 9 | Control | 59 | M | 6 |
| 10 | Control | 61 | F | 7 |
| 11 | Control | 61 | G | NA |
| 12 | Control | 70 | G | 10 |
| 13 | Control | 71 | M | 7 |
| 14 | Control | 83 | M | 12 |
| 15 | Control | 86 | M | 1.5 |

**Data S1. (separate file)**

Excel file containing source data for all figures, tables, and supplementary figures.
